# Integrating biodiversity assessments into local conservation planning: the importance of assessing suitable data sources

**DOI:** 10.1101/2023.05.09.539999

**Authors:** Thibaut Ferraille, Christian Kerbiriou, Charlotte Bigard, Fabien Claireau, John D. Thompson

## Abstract

Strategic Environmental Assessment (SEA) of land-use planning is a fundamental tool to minimize environmental impacts of artificialization. In this context, Systematic Conservation Planning (SCP) tools based on Species Distribution Models (SDM) are frequently used for the elaboration of spatially exhaustive biodiversity diagnostics. Despite the paradigm of “garbage in - garbage out” that emphasises the importance of testing the suitability of data for SDM and priority conservation areas, the assessment of database sources remains relatively rare. In addition, the lack of practical recommendations for the use of open-access databases by SEA stakeholders remains a problem. The aim of this study is to explore the quality of data sources that can be used in SEA to assess priority conservation areas in SEA. The study used data for nine taxonomic groups (commonly used in inventories for environmental impact assessment) and three databases available to SEA stakeholders. Three local administrative entities in very different socio-ecological contexts were used to examine three main issues : (i) the suitability of local versus regional or country databases for assessing conservation priorities, (ii) differences among taxonomic groups or territories in terms of the suitability of databases, (iii) the importance of the quality of databases for the application of SDM to assess priority conservation areas. Our study provides several clear messages for potential users of open-access databases. First, the need for prudence in the interpretation of biodiversity maps. Second, the collection of individual databases at the country scale is necessary to complete local data and ensure the suitability of SDM in a local context. Third, a data driven approach can lead to the use of notably different species communities to identify priority conservation areas when compared to the community in the original database. Finally, we propose a workflow to guide SEA stakeholders through the process of data rationalization and use in conservation planning.

## 1 Introduction

Land-use change, in particular urban land expansion, leads to artificialization of habitats and soils and is one of the major causes of the loss of biodiversity (Maxwell et al., 2016; IPBES, 2019). The reduction and fragmentation of natural habitats leads to population declines and species extinction (Fahrig, 1997; Horváth et al., 2019; Lino et al., 2019), as well as biotic homogenization, i.e. mostly the extinction of specialist species and the introduction of exotic species, which involves an increase in genetic, taxonomic and functional similarity (Olden and Rooney, 2006; Zambrano et al., 2019).

A major tool to limit artificialization is the mitigation hierarchy used in environmental assessments studies. This approach consists of three sequential steps: “avoid” impacts, “reduce or minimize” impacts not avoided and “offset” residual impacts (Bull et al., 2016; Maron et al., 2016). However, in the current application of the mitigation hierarchy several weaknesses prevent it from achieving the goal of “No Net Loss” of biodiversity (Quétier et al., 2014; Bezombes et al., 2019). Avoidance is poorly implemented despite the fact that it is the first and most efficient step of the hierarchy (Bigard et al., 2017; Phalan et al., 2018). What is more, the mitigation hierarchy is mostly applied in a project-by-project approach without scaling up (Pope et al., 2013; Bigard et al., 2017), which limits proper consideration of fragmentation issues (Gontier et al., 2006) and cumulative impacts (Whitehead et al., 2017), including those of multiple small projects (Bigard et al., 2017).

To anticipate avoidance measures, Strategic Environmental Assessment (SEA) of land-use planning is a global and fundamental tool to minimize environmental impacts (Baker et al., 2005). SEA provides for the integration of avoidance measures early in the land-use planning process through environmental assessment of policies, plans and programs (Fundingsland Tetlow and Hanusch, 2012; Bigard et al., 2020). However, the implementation of SEA is often based on biodiversity diagnostic maps that are rarely complete and exhaustive. Indeed, biodiversity diagnostics are rarely exclusively based on empirical observations from field surveys (Phalan et al., 2018) and usually use areas and documents already known (e.g., protected areas and green infrastructures).

Spatial modelling provides a tool for the elaboration of spatially exhaustive diagnostics of biodiversity maps for land-use and conservation planning (Almenar et al., 2019; Tarabon et al., 2019; Bigard et al., 2020; Tulloch et al., 2019; Baker et al., 2021; Boileau et al., 2022). Among these methods of biodiversity modelling, Species Distribution Models (SDM) are widely used to predict suitable habitat for species based empirical observations (Guisan et al., 2017; Zurell et al., 2020) and are increasingly used in conservation planning (Guisan et al., 2017; Domisch et al., 2019; Baker et al., 2021). Systematic Conservation Planning (SCP) tools are also particular pertinent to identify priority biodiversity stakes and avoid the adoption of an *ad hoc* approach (Margules and Pressey, 2000; Pressey and Bottrill, 2008) in order to inform SEA (Tulloch et al., 2019).

The management of databases and their use for conservation planning is a critical issue for the application of such methods to practical conservation planning. The databases available for SEA stakeholders (i.e. decision makers, environmental consultants and conservation managers) are often limited because of data sensitivity or ownership issues, although more and more programs contain data that are publicly available and use of them can be made without any particular attention to their quality (Costello and Wieczorek, 2014; Tittensor et al., 2014) and they are generally unfamiliar to SEA stakeholders. Surprisingly however, despite the prevailing recognition of the “garbage in – garbage out” that emphasises the critical importance of the quality of data (Sanders and Saxe, 2017; Canbek, 2022), an examination of data suitability is relatively rare in local conservation planning (Rondinini et al., 2006; Hermoso et al., 2015a). In this context, some authors argue the necessity of examining the sensitivity of model results to the nature of the datasets that are used (Sanders and Saxe, 2017; Clare et al., 2019; Velazco et al., 2020). SDM studies generally use data that has not been designed specifically for this type of analysis, and is often comprised of presence-only data, hence the need for a rigorous assessment of sampling biases (Beck et al., 2014; Botella et al., 2018; Guisan et al., 2017). Another particularly important point that can influence distribution modelling is the spatial extent of the data, and in particular the question of whether to use only local data or those collected on a larger scale (Baker et al., 2021; Meyer, 2007). The choice and possible combination of data sources is part of this problem due to the fact that they often vary considerably in their design, the gradients covered, and potential sampling biases (Fletcher et al., 2019; Boyd et al., 2023). Basically, the use of available databases requires a rigorous test of their quality and pertinence (Zuckerberg et al., 2011), especially when used for analyses such as SDM (Tulloch et al., 2016; Domisch et al., 2019). Confidence in the models must be assessed through the use of metrics adapted to the data (Guisan et al., 2017; Leroy et al., 2018). As recognised by Clare et al., (2019), the lack of practical recommendations for the use of databases that differ in terms of their quality and pertinence by public authorities or other institutions remains a serious problem.

The overall goal of this study is to test the influence of different database sources that can be used by SEA stakeholders to map priority conservation areas in SEAs based on SCP. To do so, we studied three local administrative territories that occur in different socio-ecological contexts in France. The study has three main objectives. First, we assess the content of three open-access databases for nine taxonomic groups commonly used in naturalist inventories in environmental assessment studies. We evaluate their suitability in terms of data quantity for SDM application, at three scales (local, regional and country). SDM and SCP analyses were performed for two taxonomic groups (Aves and Papilionidae) to test the hypothesis that sampling bias and differences in ecological response scales of species may influence the identification priority conservation areas. Second, we explore the influence of databases on the application of SDM to assess priority conservation areas. Third, we analyse the influence of this data-driven approach on the composition of species communities that are ultimately used in the identification of priority conservation areas relative to the actual communities in the original databases.

## 2 Methods

### 2.1 Study sites

To assess the availability and suitability of pertinent data sources, we selected three French local administrative entities in charge of land-use planning: Lodévois-Larzac (T1), Brocéliande (T2), La Rochelle (T3). We selected these study sites on the basis their contrasting social, ecological and geographical contexts in order to examine patterns of variation of data suitability among sites (Figure 1, Table 1). For example, each of the three study sites have different ecosystems and bioclimates and the sites vary in terms of urbanization pressures from 3% to 28% artificial land-use cover for sites T1 and T2 respectively. This is due to the presence of a major city (La Rochelle) in the T2 study site. The major towns in the other two study sites are smaller however there is a major city less than 50 km away for both of them (Montpellier and Rennes respectively). Only site T1 has an important cover of protected areas (Natura 2000) area.

**Figure 1.**
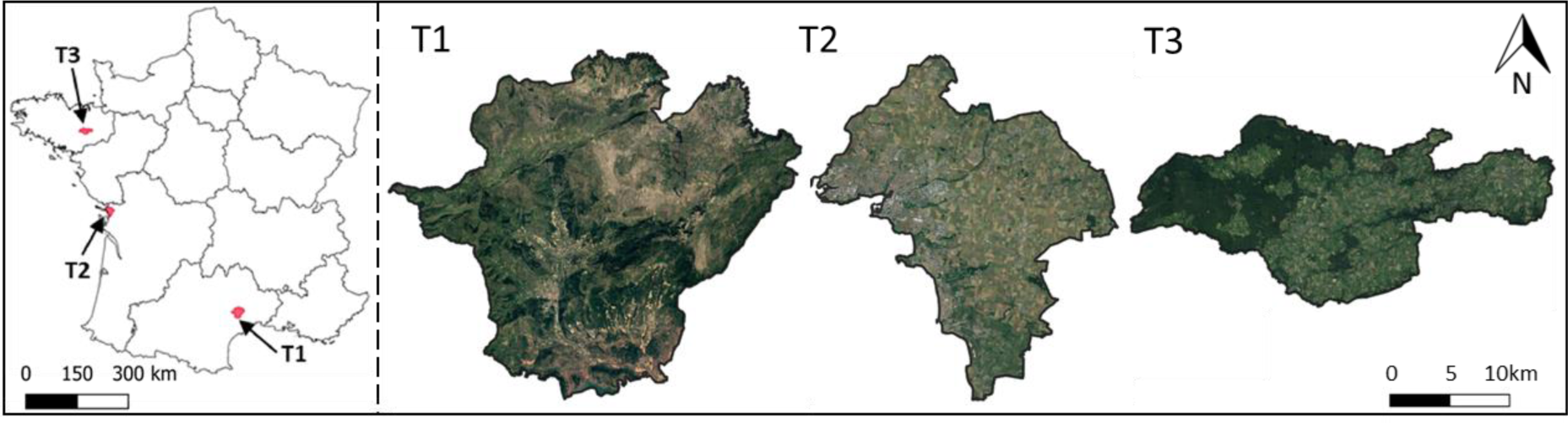
Localization of the study sites in French administrative regions: T1 is Lodévois-Larzac, T2 is La Rochelle, T3 is Brocéliande. Source: IGN, Google, 2023.

**Table 1.**
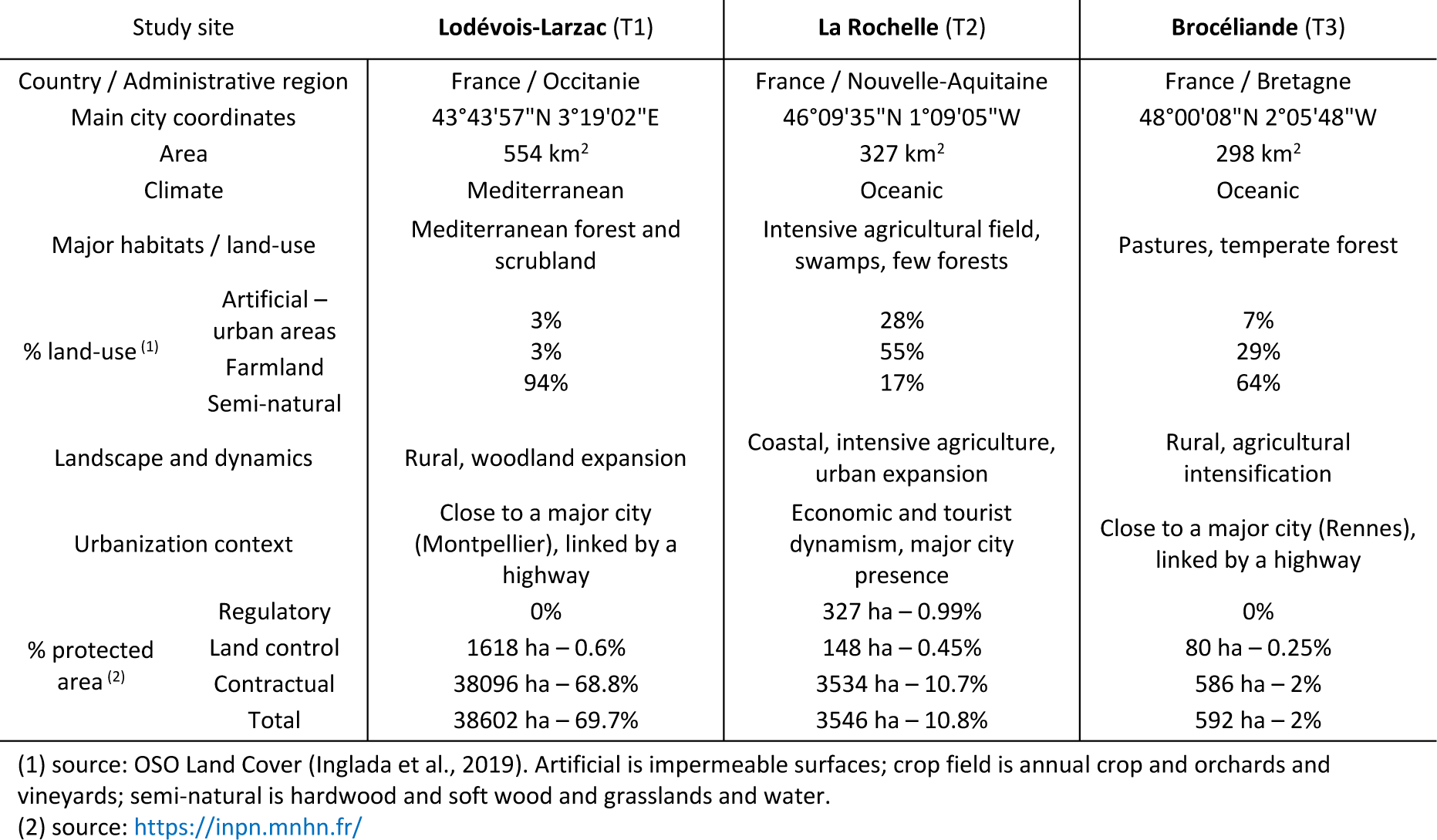
Description of three study sites in France.

### 2.2 Workflow of analysis

A methodological framework was developed to test the influence of different database sources in mapping priority conservation areas in SEAs thanks to a SCP approach, all steps are summarized in Figure 2 and in the following text.

**Figure 2.**
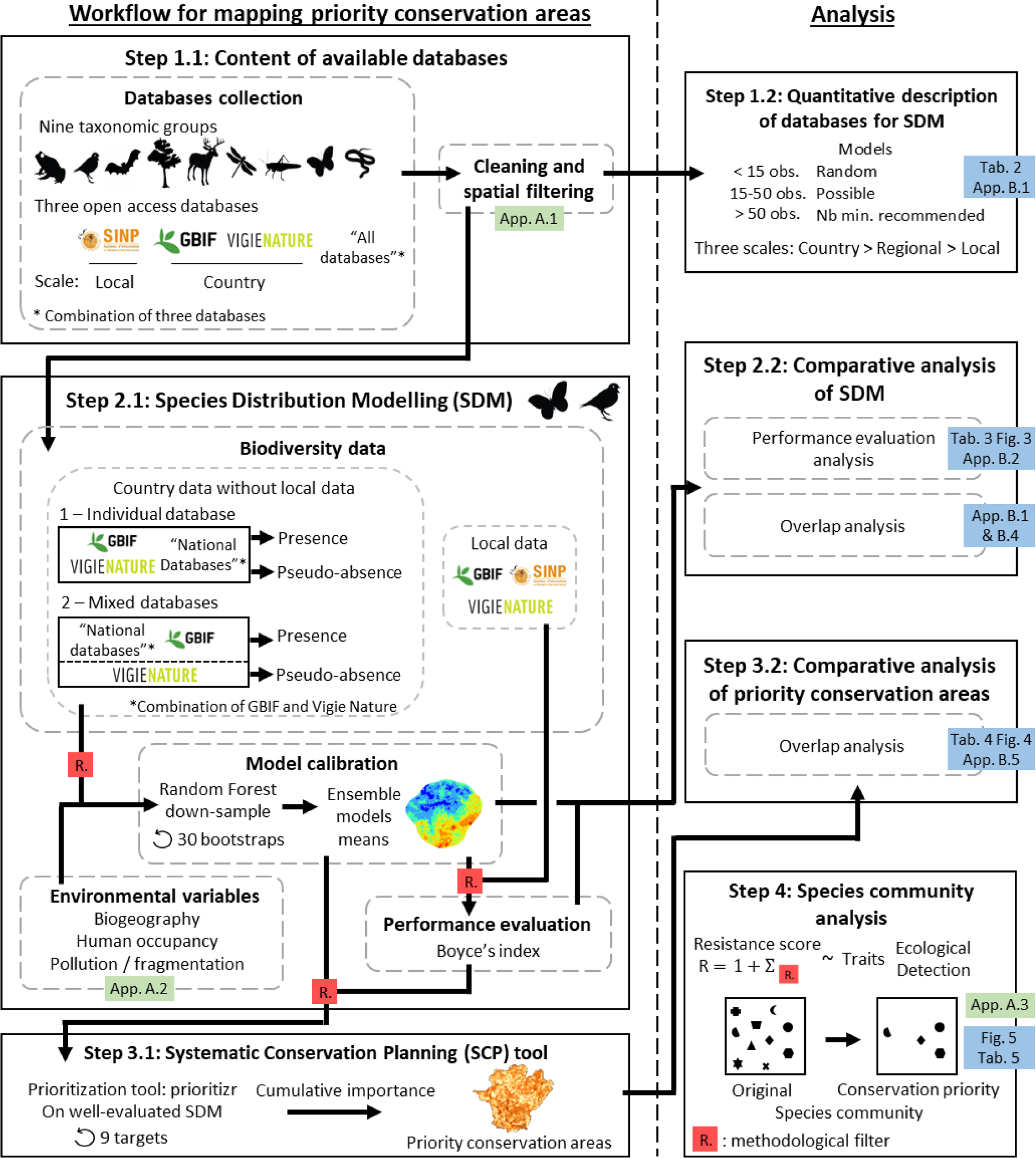
Methodological framework applied to test the influence of different database sources to map conservation planning areas in SEAs based on SCP. The green and blue boxes are the appendices detailing the methodology and the results of the analyses, respectively. The red boxes “R.” are the methodological filters used for generate workflow resistance score.

### 2.3 Databases available for SEA stakeholders

#### 2.3.1 Content of available databases

We focused on nine taxonomic groups commonly used for naturalist inventories for environmental impact assessment studies: Amphibia, nesting Aves (hereafter name Aves), Chiroptera, Flora, Mammalia aptera (hereafter name Mammalia), Orthoptera, Odonata, Papilionidae and Reptilia (Bigard et al., 2017; Guillet et al., 2019; Iorio et al., 2022). The three spatial scales used for data collection are depicted in Figure 1: the local scale (i.e. study site with a 10km buffer around), the regional scale (i.e. French administrative regions); and the country scale (i.e. continental France). We selected three open-access databases containing these groups that can be widely used by SEA stakeholders for the assessment and hierarchy of conservation priorities (Figure 2, step 1.1). This study thus directly addresses SEA stakeholders (i.e. decision makers, environmental consultants and conservation managers) using the databases available to them.

The first of these databases concerns the French Natural and Landscape Information System (SINP) that is structured at the scale of French administrative regions in charge of data extraction requests. Each site has its own database, can be collected only at local scale due to the limited extent of data requests, without the need for a special request (maximum 2000 km^2^, i.e. nearby 10km buffer zone around the study site). This database is composed of opportunist observations and only contains presence data for taxa for which identification is confirmed by experts (Jomier et al., 2018) (see Appendix A.1).

The second database is the Global Biodiversity Information Facility (GBIF) an international platform for the provision of biodiversity data that is based on information collected from various databases (Telenius, 2011). It is composed of observation data that are not based on protocols and for which presence data and identification are not subject to expert confirmation. The data downloading is autonomous from the website (see Appendix A.1).

The third database is a French biodiversity monitoring scheme (Vigie Nature) dedicated to assess spatio- temporal populations trends. Within this monitoring scheme data collection is based on a standardized biodiversity survey. Despite local spatial distribution heterogeneities, the sampling plan ensures a representation of the current distribution of habitats and landscapes across France (Julliard and Jiguet, 2002; Mariton et al., 2022). Homogeneity in identification criteria and compliance with the protocol are ensured by offering training to volunteers. This database is composed of presence/absence and abundance data. Access to these databases requires a data extraction request to the person in charge (see Appendix A.1).

These three databases were combined in two ways: “All databases” (i.e. the combination of SINP, GBIF and Vigie Nature) used in section 2.3 (i.e. for assess which taxa are enough documented within each dataset) and “National databases” (i.e. the combination of GBIF and Vigie Nature, which are available at country scale) used specifically in section 2.4 (i.e. to test the effect of database sources on SDM performance).

The databases were collected for continental France except for the SINP that was collected in 10km buffer zone around the study sites due to the access restrictions explained above. The databases were collected over a period of 11 years (i.e. from 01/01/2010 to 31/12/2020) and data with spatial inaccuracy greater than 50 meters were not considered. We made a series of operations to standardize, correct and homogenize taxa names at the specific taxonomic level using the French taxonomic reference “TAXREF.V14” (Gargominy et al., 2021). We transformed data into occurrences and limited their sampling biases by geographical filtering using “spThin” package (Aiello-Lammens et al., 2015). In each study site, we identify the presence of one species at least five observations from “All databases” combined at local scale and defined as present species in France by TAXREF.V14 (Appendix A.1).

#### 2.3.2 Quantitative description of databases for SDM

For the nine taxonomic groups, four metrics were selected to quantitatively describe the amount and thus the suitability of each database (i.e. SINP, GBIF, Vigie Nature and “All databases”) for the realization of SDM in presence-only (Figure 2, step 1.2): (i) the number of species observed in the study site; (ii) the proportion of species with < 15 observations which represents the minimal threshold for the utility of SDM with more accurate predictions than in a random model (Støa et al., 2019); (iii) the proportion of species with between 15 and 50 observations, i.e. the minimum number of presences recommended for SDM (Merow et al., 2014 in Guisan et al., 2017); (iv) the proportion of species with > 50 observations, i.e. highly suitable for modelling (Støa et al., 2019). These metrics were calculated at three different scales: (a) for each study site including a 10km buffer zone (local scale) that is the maximum extent for a SINP data request; (b) on a regional scale that is used for the structure of biodiversity data in France; (c) for continental France. SINP database is analysed individually only at the local scale due to the previously mentioned restriction of access, nevertheless it integrated the three scales of the “All databases”.

### 2.4 Systematic conservation planning process

For the three study sites, we identified priority conservation areas with a Systematic Conservation Planning (SCP) tool based on Species Distribution Models (SDM) (Figure 2, steps 2.1 and 2.2). Several variants of SDM were made using different database sources (i.e. GBIF, Vigie Nature and “National databases”) and two methods of generating pseudo-absences (named individual database or mixed databases). Among the nine taxonomic groups studied above, only Aves and Papilionidae taxa were analysed to compare the tests in the SCP process. The data available in France for these groups is sufficient in quantity to realize SDM with each database. The use of these two groups allows for a comparison between one group of highly mobile taxa with a large home range (Aves) and another group with a smaller home range and whose movement closely tracks local environmental variation (Papilionidae). These two taxonomic groups thus have different biological traits associated with their dispersal and function, hence we predict differences in in terms of the spatial resolution of their distribution.

#### 2.4.1 Species distribution modelling (SDM)

We modelled favourable habitats for birds and butterflies in the three study sites using SDM (Figure 2, step 2.1). A resolution of 50m was used to meet the needs of the SEA of land-use planning. A buffer zone of 10km around each of the study sites (i.e. local scale) was used for the SDM prediction to limit any border edge effects and to increase the number of species that could be modelled and evaluated. Indeed, species with less than 15 data points for the calibration (threshold explain above, Støa et al., 2019) and/or less than 10 data for performance evaluation (threshold defined by expert opinion) were not modelling.

Biodiversity data used for the SDM came from the databases described above at the country scale, according to the results of section 2.3 (Table 2). These data were separated into two independent datasets that allow for robust validations with independent data (Matutini et al., 2021). Country data without local data were used for model calibration and local data were used only for model performance evaluation. Therefore, the SINP database, which is only available at local scale, was not used for model calibration.

**Table 2.** Quantitative description of the observation databases (Local SINP, GBIF, Vigie Nature) for nine taxonomic groups in the three study sites (“T1”: Lodévois- Larzac, “T2”: La Rochelle, “T3”: Brocéliande) for “local” (study site with buffer of 10km), “Regional” (administrative region) and “Country” (continental France). “nSp” is the number of species observed at least 3 times in a given study site within a 10 km buffer zone, “>15obs” is the percentage of species with more than 15 observation and “>50obs” the percentage with more than 50 observations (red < 25%, orange 25-75%, green >75%, grey - no data).

| Taxa |  | Local |  |  | GBIF<br>Regional |  |  | Country |  |  | SINP<br>Local |  |  | Local |  |  | Vigie Nature<br>Regional |  |  | Country |  |  | All databases combined |  |  |  |  |  |  |  |  |
| --- | --- | --- | --- | --- | --- | --- | --- | --- | --- | --- | --- | --- | --- | --- | --- | --- | --- | --- | --- | --- | --- | --- | --- | --- | --- | --- | --- | --- | --- | --- | --- |
|  |  | T1 | T2 | T3 | T1 | T2 | T3 | T1 | T2 | T3 | T1 | T2 | T3 | T1 | T2 | T3 | T1 | T2 | T3 | T1 | T2 | T3 | T1 | T2 | T3 | T1 | T2 | T3 | T1 | T2 | T3 |
| Amp | nSp | 11 | 1 | 11 | 13 | 2 | 11 | 14 | 2 | 11 | 13 | 2 | 0 | No program |  |  |  |  |  |  |  |  | 14 | 2 | 11 | 14 | 2 | 11 | 14 | 2 | 11 |
|  | >15obs |  |  |  |  |  |  |  |  |  |  |  |  |  |  |  |  |  |  |  |  |  |  |  |  |  |  |  |  |  |  |
|  | >50obs |  |  |  |  |  |  |  |  |  |  |  |  |  |  |  |  |  |  |  |  |  |  |  |  |  |  |  |  |  |  |
| Ave | nSp | 100 | 73 | 77 | 198 | 159 | 120 | 204 | 170 | 131 | 197 | 94 | 82 | 114 | 155 | 101 | 182 | 143 | 120 | 201 | 170 | 132 | 205 | 171 | 132 | 205 | 166 | 132 | 205 | 171 | 132 |
|  | >15obs |  |  |  |  |  |  |  |  |  |  |  |  |  |  |  |  |  |  |  |  |  |  |  |  |  |  |  |  |  |  |
|  | >50obs |  |  |  |  |  |  |  |  |  |  |  |  |  |  |  |  |  |  |  |  |  |  |  |  |  |  |  |  |  |  |
| Chi | nSp | 4 | 0 | 5 | 13 | 13 | 5 | 19 | 19 | 21 | 21 | 6 | 18 | 25 | 22 | 24 | 25 | 22 | 24 | 25 | 22 | 25 | 26 | 22 | 26 | 26 | 22 | 26 | 26 | 22 | 26 |
|  | >15obs |  |  |  |  |  |  |  |  |  |  |  |  |  |  |  |  |  |  |  |  |  |  |  |  |  |  |  |  |  |  |
|  | >50obs |  |  |  |  |  |  |  |  |  |  |  |  |  |  |  |  |  |  |  |  |  |  |  |  |  |  |  |  |  |  |
| Flo | nSp | 786 | 476 | 639 | 1428 | 1093 | 825 | 1573 | 1261 | 995 | 1703 | 1335 | 843 | 286 | 177 | 136 | 825 | 657 | 413 | 1336 | 1105 | 848 | 1844 | 1460 | 1093 | 1844 | 1158 | 1093 | 1844 | 1460 | 1093 |
|  | >15obs |  |  |  |  |  |  |  |  |  |  |  |  |  |  |  |  |  |  |  |  |  |  |  |  |  |  |  |  |  |  |
|  | >50obs |  |  |  |  |  |  |  |  |  |  |  |  |  |  |  |  |  |  |  |  |  |  |  |  |  |  |  |  |  |  |
| Mam | nSp | 10 | 1 | 14 | 23 | 6 | 21 | 23 | 6 | 28 | 23 | 7 | 30 | No program |  |  |  |  |  |  |  |  | 23 | 7 | 30 | 23 | 6 | 30 | 23 | 7 | 30 |
|  | >15obs |  |  |  |  |  |  |  |  |  |  |  |  |  |  |  |  |  |  |  |  |  |  |  |  |  |  |  |  |  |  |
|  | >50obs |  |  |  |  |  |  |  |  |  |  |  |  |  |  |  |  |  |  |  |  |  |  |  |  |  |  |  |  |  |  |
| Odo | nSp | 38 | 5 | 23 | 56 | 7 | 28 | 57 | 7 | 30 | 57 | 1 | 0 | 0 | 1 | 24 | 56 | 7 | 27 | 57 | 7 | 30 | 57 | 7 | 30 | 57 | 7 | 30 | 57 | 7 | 30 |
|  | >15obs |  |  |  |  |  |  |  |  |  |  |  |  |  |  |  |  |  |  |  |  |  |  |  |  |  |  |  |  |  |  |
|  | >50obs |  |  |  |  |  |  |  |  |  |  |  |  |  |  |  |  |  |  |  |  |  |  |  |  |  |  |  |  |  |  |
| Ort | nSp | 53 | 3 | 24 | 93 | 25 | 26 | 95 | 25 | 34 | 91 | 8 | 0 | 25 | 19 | 19 | 25 | 19 | 19 | 25 | 19 | 19 | 98 | 25 | 34 | 98 | 25 | 34 | 98 | 25 | 34 |
|  | >15obs |  |  |  |  |  |  |  |  |  |  |  |  |  |  |  |  |  |  |  |  |  |  |  |  |  |  |  |  |  |  |
|  | >50obs |  |  |  |  |  |  |  |  |  |  |  |  |  |  |  |  |  |  |  |  |  |  |  |  |  |  |  |  |  |  |
| Pap | nSp | 115 | 15 | 33 | 144 | 18 | 43 | 147 | 18 | 52 | 154 | 11 | 0 | 0 | 0 | 50 | 132 | 18 | 51 | 149 | 18 | 52 | 154 | 18 | 52 | 154 | 18 | 52 | 154 | 18 | 52 |
|  | >15obs |  |  |  |  |  |  |  |  |  |  |  |  |  |  |  |  |  |  |  |  |  |  |  |  |  |  |  |  |  |  |
|  | >50obs |  |  |  |  |  |  |  |  |  |  |  |  |  |  |  |  |  |  |  |  |  |  |  |  |  |  |  |  |  |  |
| Rep | nSp | 18 | 2 | 7 | 20 | 4 | 7 | 21 | 4 | 7 | 21 | 4 | 0 | No program |  |  |  |  |  |  |  |  | 22 | 4 | 7 | 22 | 4 | 7 | 22 | 4 | 7 |
|  | >15obs |  |  |  |  |  |  |  |  |  |  |  |  |  |  |  |  |  |  |  |  |  |  |  |  |  |  |  |  |  |  |
|  | >50obs |  |  |  |  |  |  |  |  |  |  |  |  |  |  |  |  |  |  |  |  |  |  |  |  |  |  |  |  |  |  |

For model calibration, pseudo-absences were generated with two methods, separately for each taxonomic group. First, methods to generate pseudo-absences in the individual databases for all their data (i.e. GBIF, Vigie Nature and “National databases”). For databases with a protocol for sampling (i.e. Vigie Nature) to optimize species detection (day and year periods), the absence points were defined as all the points without the observed species. For databases without such sampling protocols for all their data (i.e. GBIF and “National databases”), pseudo-absence data were generated with the target-group (TG) approach, which infers the sampling bias from the aggregated occurrences of (TG) species, i.e. the respective taxonomic groups (Ponder et al., 2001; Anderson, 2003; Phillips et al., 2009). Second, a method to mix the presence data in the GBIF and “National databases” with the absence from Vigie Nature (named mixed databases) was applied (Hermoso et al., 2015a).

Three types of environmental variables were used for SDM: geographic, human occupancy and pollution and fragmentation (Appendix A.2).

SDM were calibrated by Random Forest down-sample (Valavi et al., 2021a) which according to Valavi et al. (2021b) is among the best performing models for presence-only data. Although the Random Forest is not very sensitive to the non-independence of the variables and over-parametrization (Matsuki et al., 2016; Srisa-An, 2021), in order to be parsimonious, the collinear variables were removed (Pearson >0.7, Appendix A.2, Brun et al., 2020). Thirty bootstraps were performed for each SDM (Guisan et al., 2017) using a calibration for 70% of the data at the country scale outside of local scale. The thirty Random Forest bootstraps were combined with mean to provide an ensemble prediction of habitat suitability for all species.

The performance evaluation of the models was done using the Boyce’s index (CBI), the most suitable metric for model in presence-only (Boyce et al., 2002; Leroy et al., 2018), with local data (i.e. “All databases” combining SINP, GBIF and Vigie Nature). Dubos et al. (2022) reveal the CBI turns out to be misleading in some cases, thus we used a threshold of 0.3 to define good or poor model quality.

#### 2.4.2 Systematic conservation planning (SCP) tool

Priority conservation areas in the three study sites were analysed from SDMs for each database source in using a SCP tool (Figure 2, step 3.1). To meet the needs of SEAs, the study site was restricted to administrative boundaries with a buffer zone of 1km to maintain coherence between administrative entities and a resolution of 50m. The aim of SEA biodiversity conservation strategies is to establish priorities for the whole study site as a whole and all the cells have the same cost value of 1. The objective was a maximum coverage objective that seeks to maximize the number of features, i.e. the SDMs (Church et al., 1996). The features were only the good quality models defined previously. The priority conservation areas decisions were between 0 and 1. To obtain a priority gradient, we cumulated Ferrier importance scores (Ferrier et al., 2000) from nine targets of the total amount of each feature (from 0.1 to 0.9 every 0.1). We used the package “prioritizr” (Hanson et al., 2021) with the open-source solver SYMPHONY (Kim et al., 2023).

### 2.5 Comparative analysis of SDM and priority conservation areas

We analysed the influence of database sources on SDM predictions and priority conservation areas (Figure 2, step 2.2 and 3.2). SDM performance evaluations were analysed between database sources. The SDM prediction and priority conservation areas maps were compared with the Spearman’s rank coefficient (Phillips et al., 2009) and the Schoener’s D index as a measure of projection overlap (Schoener, 1968) which was calculated with the ENMTool R package (Warren et al., 2008; Warren and Dinnage, 2022).

### 2.6 Species community analysis

We assessed the influence of the complete data driven workflow on the composition of species communities, i.e. differences between the original community (i.e. all species observed in study site) in the database and the final community used to identify priority conservation areas (Figure 2, step 4).

To do so, we developed a workflow resistance score for each of the methodological filters for all species. A score of 1 is allocated to species observed in the study area that did not cross any of the stepwise filters. A score of 2 is allocated to species with sufficient data to calibrate SDM, i.e. > 15 country observations, or to evaluate the performance of SDM, i.e. > 10 local observations. A score of 3 is allocated to species with sufficient data to calibrate and evaluate the performance of the SDM. A score of 4 is for species that were present in the final analysis as a priority species for conservation planning (i.e. with the two previous filters and a Boyce’s index > 0.3). In order to assess species composition bias ultimately considered in priority conservation areas, species communities were analysed through traits that can influence species detection (mass, displacement capacity, period activity) and ecological traits (habitats, specialisation) (Appendix A.3). Missing data were completed with a trait imputation procedure generated using the R package “missForest” (Stekhoven and Bühlmann, 2012) by considering evolutionary relationships in the imputation process (see Carmona et al., 2021) using the R script of Toussaint et al. (2021). Due to the nature of the response variable (i.e. ordinal scoring including four modalities), we used ordinal regression mixed models with cumulative link using the clmm function of “ordinal” R package (Christensen, 2022). We adapted the link function to the data distribution, using a “cauchit” link for Aves and a “logit” link for Papilionidae. Species traits were used as fixed effects, while the random effects selected were the study sites for Aves and the combination for study sites and database sources for Papilionidae. Finally, we evaluated the quality of the full model by comparing to the null model with Akaike’s information criterion (AIC) (Mac Nally et al., 2018).

Thus, our models were structured in the following way:

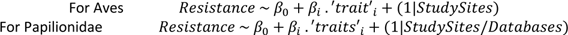

Where β is the parameter estimates, i correspond to the variables of ‘traits’ using in fix effect, 0 is the shift between ordinal class of resistance (i.e. 1|2, 2|3 and 3|4) and “1|” is the random effect.

## 3 Results

### 3.1 Quantitative description of databases for SDM

The use of individual observation databases provided a limited number of observation data for distribution modelling of many species, regardless of the taxonomic group (Table 2, Appendix B.1). None of the GBIF and SINP individual databases have more than 50 observations for all nine taxonomic groups, and there are currently no programs for three of the nine taxonomic groups in the Vigie Nature databases. Local databases and, to a lesser extent, regional databases are not equivalent in terms of the amount of data available for different taxa for the three study sites. The difference is particularly pronounced for the SINP databases, with no data available for five taxonomic groups in the T3 study site, whereas in the T1 study site, six taxonomic groups have sufficient data to model over 50% of the species. At the regional scale, the number of observations per species is highly variable among study sites for the GBIF databases and is more similar for the Vigie Nature databases.

The use of combined databases (i.e. “All databases”) increased the total number of species that can be studied. At the local scale, the proportion of species in each suitability class showed only small changes, while at the country scale of France, it allowed a significant gain in species with suitable data (Table 2). Indeed, on a country scale, the GBIF and Vigie Nature databases are complementary with each other. For example, the GBIF has few Chiroptera data, which is complemented by Vigie Nature data, and vice versa for Amphibian data.

At the country scale, aggregation of the databases seem to provide the most suitable setup (databases and scale) for SDM analysis. Using these compiled, country databases provides a large amount of data for a large number of species present in the three study sites (Table 2 and Appendix B.1).

### 3.2 SDM and priority conservation areas analysis

The evaluation of SDM revealed differences among the database sources; none of which produced more than 87% of satisfactory models for the two studied taxonomic groups and some had less than 20% of satisfactory models (Table 3, Appendix B.2). Use of the GBIF data led to a higher proportion of well-evaluated SDM, ranging between 48 and 79% of satisfactory models for the species in the two taxonomic groups. GBIF data are also more suitable than Vigie Nature data, they produced between 11% and 37% more satisfactory models than the latter database (Table 3). Nevertheless, between 4% and 9% of species provide well-evaluated models from Vigie Nature and poorly evaluated by GBIF database. The combination of “National databases” (i.e. GBIF and Vigie Nature) decreased the performance of SDMs with GBIF data, but still yield better results than SDM based on the Vigie Nature database. The substitution in individual databases of pseudo-absences for the absences from Vigie Nature (i.e. mixed databases) reduced the performance of models based on GBIF, but increased the performance of the combined “National databases” (Figure 3, Appendix B.2). Regardless of the database used, our analyses revealed significant differences between study sites (Table 3). No SDMs for butterflies in T2 could be evaluated due to insufficient local data. For the T3 study site, over 50% of the Papilionidae and Aves models perform poorly, whereas for the T1 study site poor models occur in less than 50% of the evaluations.

**Figure 3.**
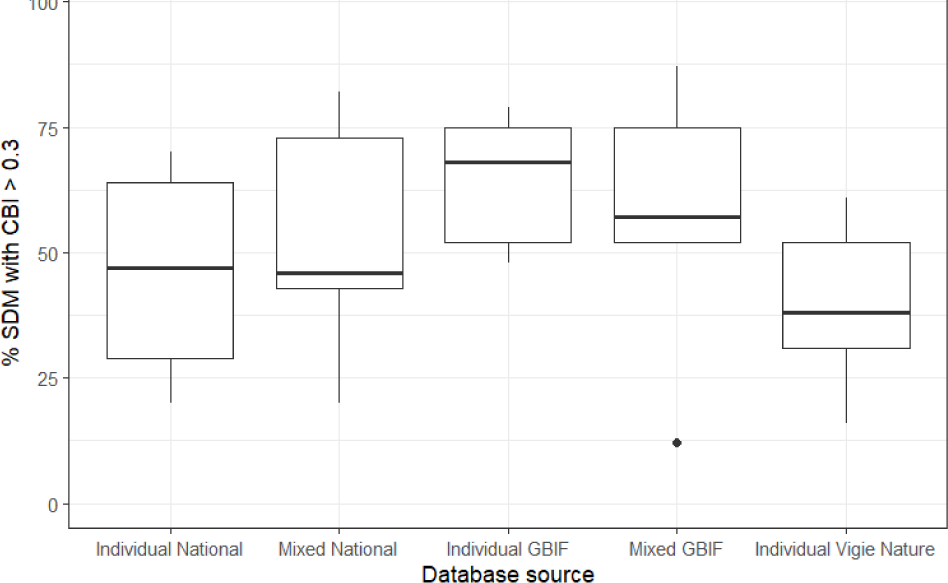
Proportion of SDM with Boyce’s index (CBI) greater than 0.3 by database source (“Individual” and “Mixed” with Vigie Nature absence), by combining study sites (T1, T2, T3) and taxonomic groups (Aves, Papilionidae). “National” combines GBIF and Vigie Nature databases.

**Table 3.** Proportion of well-evaluated SDM corresponding to Boyce’s index greater than 0.3. “National” combines GBIF and Vigie Nature databases.

| Taxa | Study site | Number of species | Individual database |  |  | SDM with Vigie Nature absence |  |
| --- | --- | --- | --- | --- | --- | --- | --- |
|  |  |  | GBIF | Vigie Nature | “National” | GBIF | “National” |
| Aves | T1 | 114 | 79 % | 61 % | 70 % | 87 % | 82 % |
|  | T2 | 87 | 68 % | 31 % | 47 % | 57 % | 46 % |
|  | T3 | 56 | 48 % | 38 % | 29 % | 52 % | 43 % |
| Papilionidae | T1 | 127 | 75 % | 52 % | 64 % | 75 % | 73 % |
|  | T3 | 25 | 52 % | 16 % | 20 % | 12 % | 20 % |

Although important differences in model performance between the database sources used for SDM showed a high degree of overlap, as indicated by Shoener’s D index with values above 0.8, the ranking of habitat suitability was highly variable. This was in particularly the case for the GBIF and Vigie Nature databases that had median spearman’s rank coefficient values between 0.2 and 0.5 and a very wide distribution (Appendix B.3). The substitution of pseudo-absence data in GBIF and “National databases” by absence data from Vigie Nature, showed a similar situation (Appendix B.4).

For priority conservation areas, whatever the individual databases used, the overlaps with Schoener’s D index were above 0.72 and similar in each study site. Nevertheless, Sperman’s rank coefficients showed a greater difference in prioritization ranks in particular between GBIF and Vigie Nature and for the T3 study site (Table 4). Between maps of priority conservation areas, we observed similarities in overlap, despite a significant difference in the hierarchy of areas to be prioritized (Figure 4, Appendix B.5). The list of species is presented in Appendix B.7, where it can be seen that there are no difficulties with respect to invasive species which are very few in the data sets.

**Figure 4.**
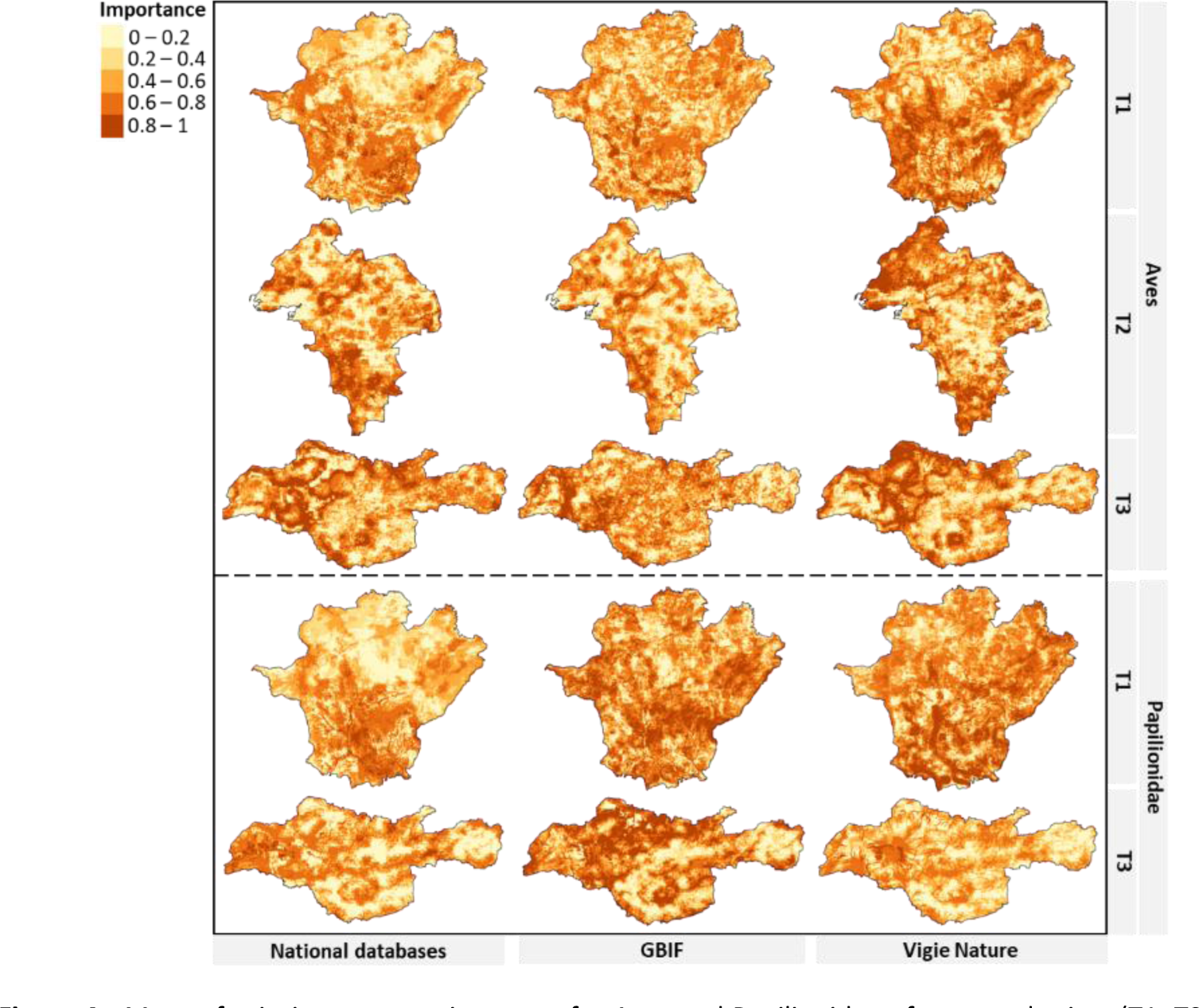
Maps of priority conservation areas for Aves and Papilionidae of tree study sites (T1, T2, T3) from different individual database source (All, GBIF, Vigie Nature).

**Table 4.** Overlap of priority conservation areas between database sources using two metrics: Schoener’s D index (D) and Spearman’s rank coefficient (S cor). VN is Vigie Nature database and “National” combines GBIF and Vigie Nature databases.

| Taxa | Study sites | Individual database |  |  |  |  |  | Pseudo-absence of individual database - absence from VN |  |  |  |
| --- | --- | --- | --- | --- | --- | --- | --- | --- | --- | --- | --- |
|  |  | "National" - GBIF |  | "National" - VN |  | GBIF - VN |  | GBIF |  | "National" |  |
|  |  | D | S cor | D | S cor | D | S cor | D | S cor | D | S cor |
| Aves | T1 | 0.82 | 0.56 | 0.81 | 0.53 | 0.78 | 0.36 | 0.75 | 0.88 | 0.79 | 0.38 |
|  | T2 | 0.82 | 0.74 | 0.76 | 0.49 | 0.72 | 0.42 | 0.76 | 0.64 | 0.79 | 0.64 |
|  | T3 | 0.77 | 0.34 | 0.81 | 0.51 | 0.78 | 0.29 | 0.73 | 0.19 | 0.79 | 0.39 |
| Papilionidae | T1 | 0.85 | 0.64 | 0.84 | 0.65 | 0.83 | 0.59 | 0.72 | 0.66 | 0.82 | 0.66 |
|  | T3 | 0.80 | 0.49 | 0.77 | 0.31 | 0.78 | 0.22 | 0.76 | 0.17 | 0.72 | 0.06 |

### 3.3 Species community analysis

The distribution of workflow resistance scores showed that only 30% and 42% respectively of Aves and Papilionidae species were integrated in priority conservation areas maps. Among species not integrated, the workflow steps filtering the most species concern the amount of suitable data for model evaluation followed by the quality of the models (Figure 5). The analysis of the species community composition observed in each of the three study sites in comparison with the species community integrated in priority conservation area identification revealed significant differences for all three study sites. For Aves communities, the differences concern an under-representation of nocturnal species, large species with high dispersal capacity, and species of swamp habitats and deciduous forests in relation to the observed species community in the databases for the three study sites. Conversely, species that favour urban habitats, shrubland, grassland and coniferous forests are over-represented in the species community of the final maps, as are species with specialized diets and foraging strata (Table 5). For Papilionidae communities, common species with long flight periods are over-represented in the final community used for analysis. Species related to anthropogenic and thermo/meso Mediterranean habitats, and species that use a wide range of hostplants are over-represented in relation to the original species community, while supra-Mediterranean species and those of montane environments are under-represented (Table 5).

**Figure 5.**
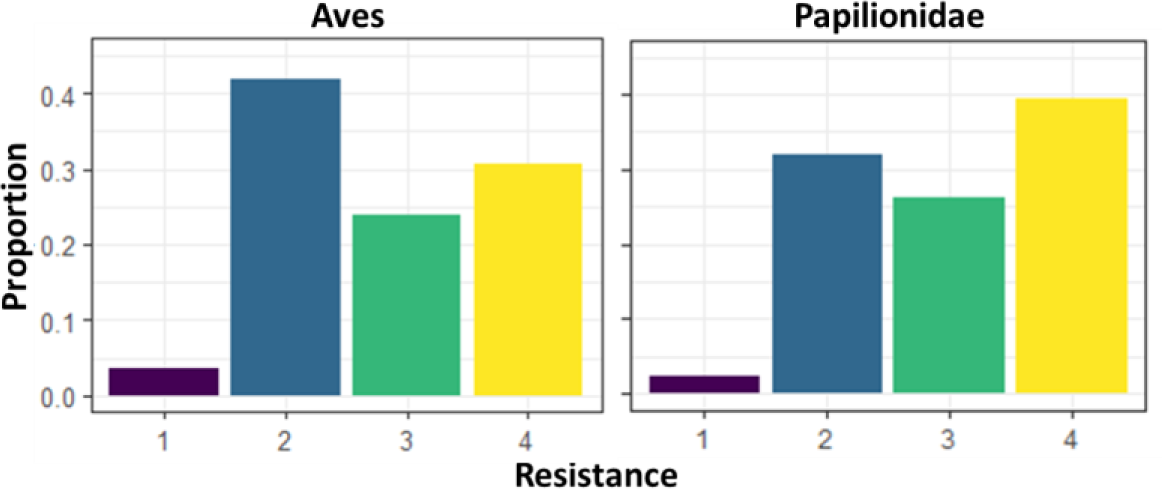
Workflow resistance scores for Papilionidae and Aves. 1 - species observed in the study area that did not cross any of the stepwise filters. 2 - species with sufficient data to calibrate SDM, i.e. > 15 country observations, or evaluate the performance of SDM, i.e. > 10 local observations. 3 - species with sufficient data to calibrate and evaluate the performance of the SDM. 4 - species present in the final analysis, i.e. Boyce’s index > 0.3.

**Table 5.** Parameter estimates (β), standard error (se) and P-values for the full model of Aves and Papilionidae species resistance to the workflow. Appendix B.6, the evaluation of the quality of the model.

|  | Variables | Aves |  |  | Variables | Papilionidae |  |  |
| --- | --- | --- | --- | --- | --- | --- | --- | --- |
| | | $\beta$ | se | P-value | | $\beta$ | se | P-value |
| Full model | $\beta_0$ 1\|2 | -7.84 | 1.12 | / | $\beta_0$ 1\|2 | -2.54 | 1.28 | / |
| | $\beta_0$ 2\|3 | 0.14 | 0.33 | / | $\beta_0$ 2\|3 | 1.39 | 1.26 | / |
| | $\beta_0$ 3\|4 | 1.4 | 0.34 | / | $\beta_0$ 3\|4 | 2.97 | 1.27 | / |
|  | Mass | -0.21 | 0.13 | . | WingspanM | 0.01 | 0.09 |  |
|  | Avian hand-wing index | -0.31 | 0.07 | *** | FMoMean | 0.38 | 0.11 | *** |
|  | Nocturn | -2.28 | 0.59 | *** | Hostplant N | 0.23 | 0.11 | * |
|  | Deciduous | -0.27 | 0.15 | . | Hostplant Spe | 0.16 | 0.11 |  |
|  | Coniferous | 0.38 | 0.15 | * | HPG Bi | 0.28 | 0.50 |  |
|  | Woodland | 0.17 | 0.14 |  | HPG Th | -0.21 | 0.22 |  |
|  | Shrub | 0.55 | 0.13 | *** | HPG Sb | -0.32 | 0.23 |  |
|  | Grassland | 0.47 | 0.13 | ** | HPG Tr | 0.22 | 0.45 |  |
|  | Mountain meadows | 0.23 | 0.24 |  | HPG Li | 0.02 | 0.36 |  |
|  | Reed | 0.08 | 0.33 |  | SSI | 0 | 0 |  |
|  | Swamps | -0.84 | 0.28 | ** | AltVeg 'A' | 0.06 | 1.12 |  |
|  | Rocks | -0.14 | 0.21 |  | AltVeg 'Mo' | -0.93 | 0.31 | ** |
|  | Urban | 0.83 | 0.14 | *** | AltVeg 'SupMed' | -0.79 | 0.46 | . |
|  | Spe. Diet | 0.92 | 0.35 | ** | AltVeg 'ThMeMed' | 1.37 | 0.37 | *** |
|  | Spe. Foraging behav. | -0.08 | 0.4 |  | Rarity '2' | 0.86 | 0.29 | ** |
|  | Spe. Diet strat | 1.49 | 0.4 | *** | Rarity '3' | 1.42 | 0.34 | *** |
|  | Spe. Habitat | -0.14 | 0.38 |  |  |  |  |  |
|  | Spe. Nest | -0.28 | 0.72 |  |  |  |  |  |
|  | Spe. Mean | -2.26 | 1.63 |  |  |  |  |  |

P-value: \*\*\* $P < 0.001$ , \*\* $P < 0.01$ , \* $P < 0.05$ , . $P < 0.1$ , / P-value not applicable
Trait description: Spe. is the specialization, FMoMean is duration of yearly flight period, Hostplant N is hostplant specificity, HostplantSpe is hostplant specificity index, HPG is hostplant growth form (Bi: short herb, Th: tall herb, Sb: shrub, Tr: tree, Li: Liana), SSI is Species Specialization Index, AltVeg is altitudinal vegetation (A=Anthropogenic, Co: Foothill, ThMeMed: Thermo/Meso-Mediterranean, Med: Mediterranean, SupMed: Supra-Mediterranean, Mo: Montane, Asa: Alpine and Subalpine). For more details on traits see Appendix A.3

## 4 Discussion

The absence of recommendations for the use of available databases that differ in terms of their quality and pertinence by public authorities or other institutions remains a serious problem for local conservation planning (Clare et al. 2019). The goal of this study was to test the suitability of different database sources that can be used by public stakeholders to map priorities for biodiversity stakes in SEAs and SCP. We found that the compilation of databases at the country scale is the most suitable procedure to apply SDM to a large number of species. For Aves and Papilionidae, the GBIF database provided the highest proportion of well-assessed SDM. We detected a significant overlap in species distributions in different database sources despite significant variability in the order of habitat suitability and similar spatial predictions for priority conservation areas. We showed that the composition of the species community used for priority conservation areas in all three study sites were clearly not representative of the observed species communities in the original database (in terms of species and ecological traits). Finally, despite important differences among the study sites in terms of the proportion of artificial land cover and protected areas we found no particular differences between the three study sites. Clearly, the data sources are the most important factor influencing the results.

Open-access biodiversity data provide a valuable source of information for decision makers, environmental consultants and conservation managers (i.e. SEA stakeholders); they contain vital information on species locations compared to expert knowledge and unshared datasets that are inaccessible to most users (Sousa-Baena et al., 2014; Meyer et al., 2015). Through their use and careful application of SDM, they contribute to the estimation of relative habitat suitability in a given study site (Baker et al., 2021). Our study showed however that SDM for a large majority of species observed locally requires their compilation on a country scale. Local and regional data are not suitable for model calibration but remain important for assessing the suitability of models in a local context. Indeed, this performance evaluation step is one of the most restrictive filters in the workflow we proposed, as evidenced by SDM’s for butterflies in the T2 study site, where no species could be evaluated. The spatial extent of data collection can influence distribution modelling (Meyer, 2007), and our study emphasises this importance for SDM and the use of data available on country scale in France.

Different types of databases are constructed in different way – with opportunistic data collection or, in some cases, as part of a scientific monitoring scheme – allowing the use of as complementary data sources (Beck et al., 2013; Shirey et al., 2021). The three individual databases examined in this study are indeed complementary in that when they are combined they provide a suitable source for modelling the distribution of many species. Nevertheless, some groups commonly have a low amount of data in such bases, e.g. Insecta (Troudet et al., 2017). The study of data with a fine spatial grain, as required for SEA and SCP (Guisan et al., 2013), reveals information gaps for more taxonomic groups at the global scale, e.g. Amphibians and Mammals (Witté and Touroult, 2017). The construction of an overall database at country scale is therefore the most appropriate way to have suitable data for SDM of different taxonomic groups. Furthermore, there is a dilemma between protocolized and opportunistic data. Although protocolized data are recommended for SDM (Guisan et al., 2017; Guillera-Arroita et al., 2015), very often the amount of such data is low, which can be detrimental at the local scale, particularly for model evaluation with data having the same sampling bias. For opportunistic data, their large number is of course a positive point, however the estimating their sampling bias can be a real challenge (Botella et al., 2018; Fithian et al., 2015; Matutini et al., 2021) to ensure the reliability of the results.

SDM of Aves and Papilionidae species clearly revealed differences between the databases used for modelling, with differential impacts on the identification of conservation priorities. Indeed, the high overlap in species distribution between data sources, as indicated by Schoener’s D index, indicates that, regardless of data source, species are predicted in similar environments (Warren et al., 2008). However, Spearman’s ranking of habitat suitability between data sources was highly variable, indicative that species’ responses to environments are highly variable, as are the location of favourable habitats (Warren et al., 2008). Although the use of presence-absence data is advocated for SDM (Guillera-Arroita et al., 2015; Valavi et al., 2021b; Dubos et al., 2022), we showed that opportunist data from GBIF provided a greater number of well-assessed models at the local scale. Models using opportunist data with a target-group approach to generate pseudo-absences provides a sufficient quality of information on species distribution (Phillips et al., 2009; Barber et al., 2022) and can be correctly used in SCP (Sofaer et al., 2019; Baker et al., 2021). The lack of data at the local scale, whatever the database, does not allow us to explain a better fit of models using GBIF data. Evaluating the models with a large proportion of opportunist data could however bias the evaluation, but only independent data were used, which provides robust validation of SDM (Matutini et al., 2021). Moreover, in contrast to Hermoso et al. (2015a), we found that mixing presence-only data with absence data increased the number of misjudged models. In addition to the use of the ROC curve (AUC) as a presence-only model evaluation metric by Hermoso et al. (2015a), the different results can be explained by different sampling biases between the two data types (Baker et al., 2022; Barber et al., 2022). Finally, the GBIF data seem to be more adapted to model the distribution of a large number of species.

The notion of “garbage in – garbage out” emphasises the critical importance of the quality of data (Sanders and Saxe, 2017), nevertheless, the examination of data suitability for conservation planning remains rare. In addition to the above issues our study revealed the importance of attention that should be paid to the representativeness of the species communities used in the models compared with the actual species communities observed in the study sites. This is particularly important in the light of the finding that there are marked differences between conservation priorities when different database sources are employed. Indeed, the number and composition of species in the community used can influence conservation priorities. Elsewhere it has been shown the difference will decline as the number of species increases (Kujala et al., 2018). The methodology tested in our study is based on a data-driven approach that attempts to use all available biodiversity data. This approach is data intensive, but is necessary to ensure the best representation of the observed local biodiversity. We revealed that such an approach can nevertheless induce a significant bias in the species community that is ultimately studied. Indeed, the prevalence of data affects the composition of the modelled species as well as the accuracy of the models and the evaluation of the species response (Fukuda and De Baets, 2016). Particular attention should thus be paid to the representativeness of the species communities used in the models in relation to the actual species communities observed in the study site.

Our study presents a workflow (Figure 2) for identifying biodiversity stakes using a data-driven approach from open-access database sources. SEA stakeholders can use this workflow as a step towards the rationalization of data in order to reduce the biases mentioned above. The confrontation of the limits of such a workflow with the needs of SEA stakeholders could illustrate how to precisely target new sources of database that should be collected according to the suitability of current databases for priority groups. This workflow could be compared with the data context of another country to compare our findings. Hermoso et al. (2015b) revealed that evaluation models using a new collection of field data does not necessarily reduce the problems of model uncertainty. However, other databases can be examined by SEA stakeholders as well as other monitoring schemes (e.g. “PopAmphibien” for Reptilian and Amphibian populations in France http://lashf.org/popamphibien-2/) or negotiate the use of databases that are not yet shared. To overcome this data sharing problem, the structuring of networks of different contributors of data and users of the databases and ambitious regional policies is necessary. As evidenced by our three study sites, the quantity of local data available is correlated with the number of years the SINP has been implemented. An important issue is thus the integration of SEA stakeholders in the workflow we propose, and their appropriation of the procedure. This could be done by a form of participatory modelling (Lagabrielle et al., 2010; Lees et al., 2021), where stakeholders are consulted for issues and choices such as the species to be examined. In such participatory modelling it is important to avoid arbitrary choices that are neither reproducible nor representative of local diversity, but rather the result of administrative or political interest. Finally, it is currently recommended to use these tools to elaborate a more holistic approach to SCP (Cadotte and Tucker, 2018).

## 5 – Perspectives: operational implementation by SEA stakeholders

Spatially exhaustive and ecologically representative priority conservation areas are crucial for the elaboration of SEAs that aim to limit artificialization impacts as early as possible in the planning process. Empirical observations are major sources of information on biodiversity that are still rarely used by SEAs. The collection of open-access databases for SEA territories provides important but incomplete knowledge on species occurrence. Furthermore, their use is particularly interesting to help strategically direct inventory campaigns (especially for under sampled taxa and areas) that go beyond the emphasis on rare, threatened and emblematic species. These additional data would clearly improve the assessment of the SDMs suitability in administrative entities as our study sites. What is also interesting here for SEA stakeholders is that the process of filtering species and attributing them a score allows for the identification of different groups of species in terms of their needs for additional data in order to undertake SDM.

The influence of database sources on the identification of priority conservation areas reveals the importance of examining their suitability. In our study this is true for three highly contrasting study areas that differ markedly in terms of the cover of protected areas and artificialisation. The problem of data sources is thus typical of many areas. Thus, it is necessary to be prudent in the interpretation of biodiversity maps. The integration of local experts may help limit any misjudgements in the workflow procedure. Indeed, the integration of “expert” knowledge and local studies is valuable information, which is important to share, and which it is important to consider in order to complete our proposal. In future studies, species conservation issues for spatial prioritization could be considered by focusing on (for example) the issues associated with threatened and/or invasive species. The multiple dimensions of biodiversity could be analyzed within a context of limited data access and the complementarity of different facets (functional and phylogenetic) in addition to a classical species-based approach (Brumm et al., 2021; Cadotte and Tucker, 2018).

A data-driven approach that considers as many species as possible requires a large amount of data, biases the species communities considered and does not highlight species of particular interest as their threats and regulatory protections. It is therefore necessary to rationalize this approach, by integrating the needs and issues of local SEA stakeholders.

## Acknowledgements

This work would not have been possible without the help of many voluntary field workers who continually provide open-access data. We are grateful to Benoit Fontaine for his work as administrator and for providing access to the Vigie Nature data, to Annegret Nicolai for collection of all open-access data on “Communauté de commune de Brocéliande (i.e. the study site “T3”), to Paul Fromage for providing access for Fauna data, to Mathieu Largarde for OEB data access, to Solène Robert for the exchanges about SINP database and Joel Kamdoum Ngueuko for access to SINP Occitanie data. We also thank Josselin Giffard and Samuel Alleaume for access to DHI NDVI, to Karine Princé for discussion on agricultural intensification variables and to Jean-Pierre Moussus for access to Papilionidae trait databases and Jeremy Froidevaux for the preview access to the EuroBaTrait https://doi.org/10.1038/s41597-023-02157-4. We thank Nicolas Dubos and Boris Leroy for discussion on the use of SDM and Eric Durand for more general discussion of this study. Finally, we also acknowledge the reviewers which improved this version.

## Data, scripts, code, and supplementary information availability

Observation data from SINP and Vigie nature databases are not available for confidentiality reasons, but their link and the request process are detailed in Appendix A1. The data of GBIF are available online: https://doi.org/10.15468/dl.ry6uw7.

The data, R scripts, outputs of steps 1.2 to 4 of figure 2 are available online: https://doi.org/10.5281/zenodo.7883973.

## Appendix

### Appendix A - Additional material and methods

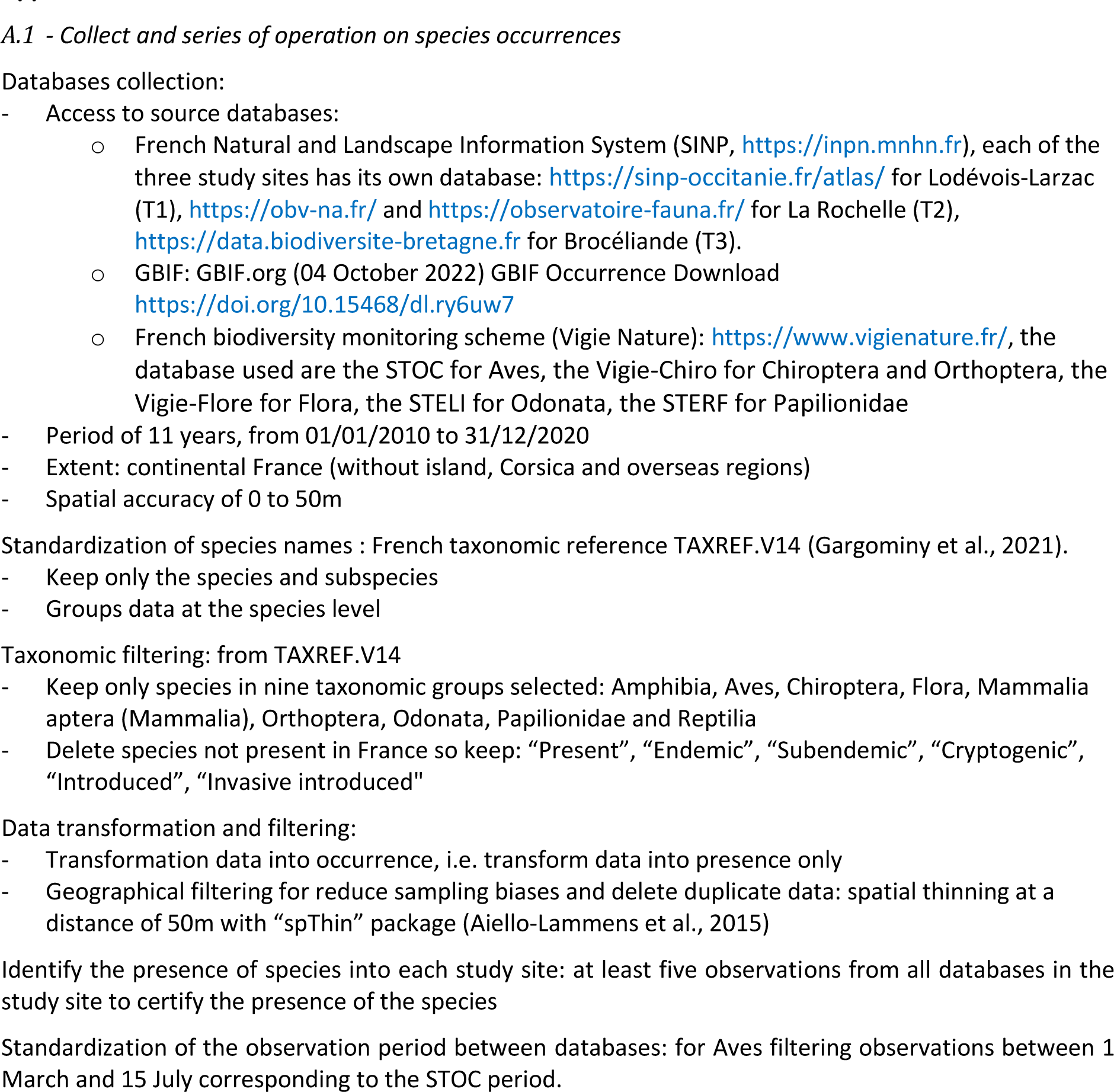
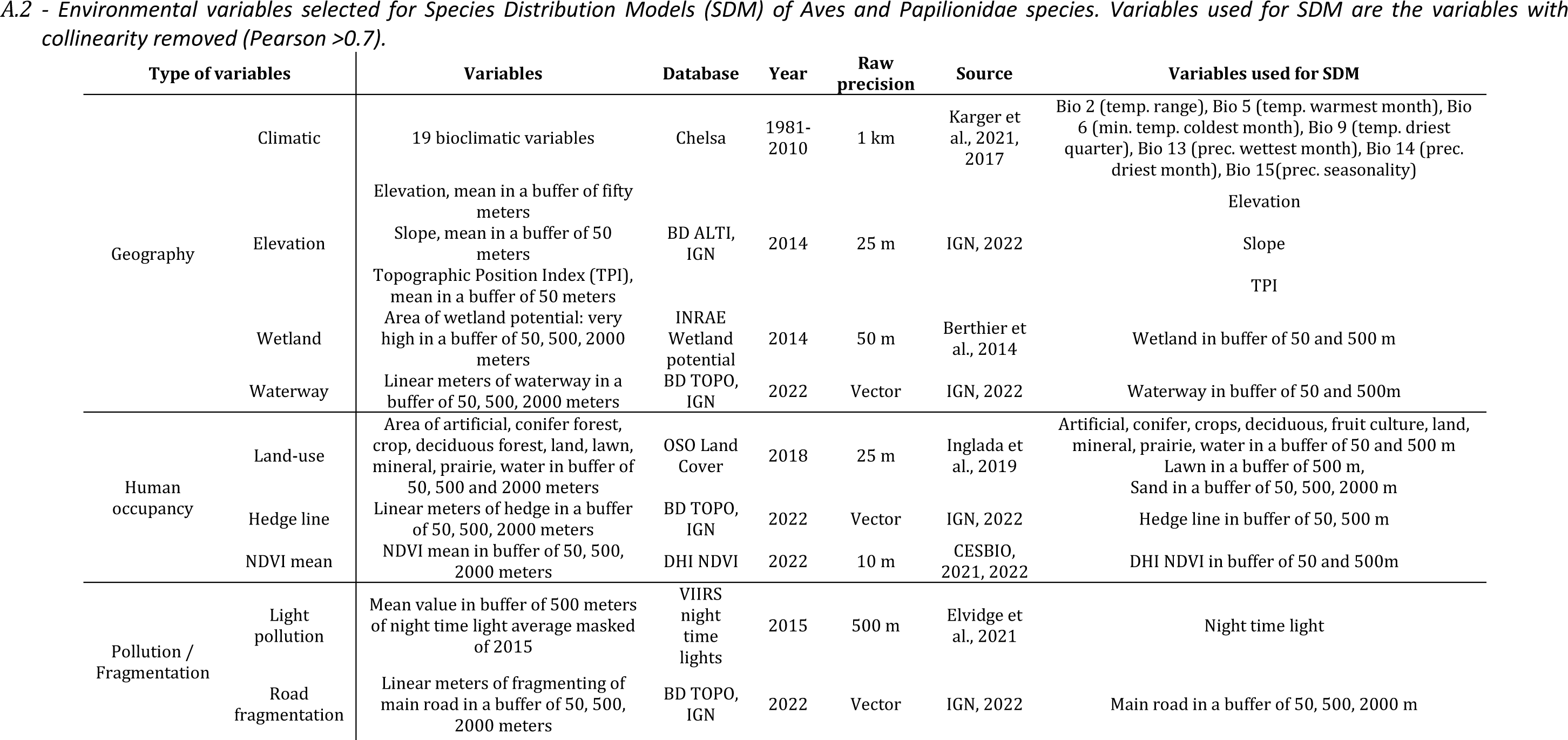
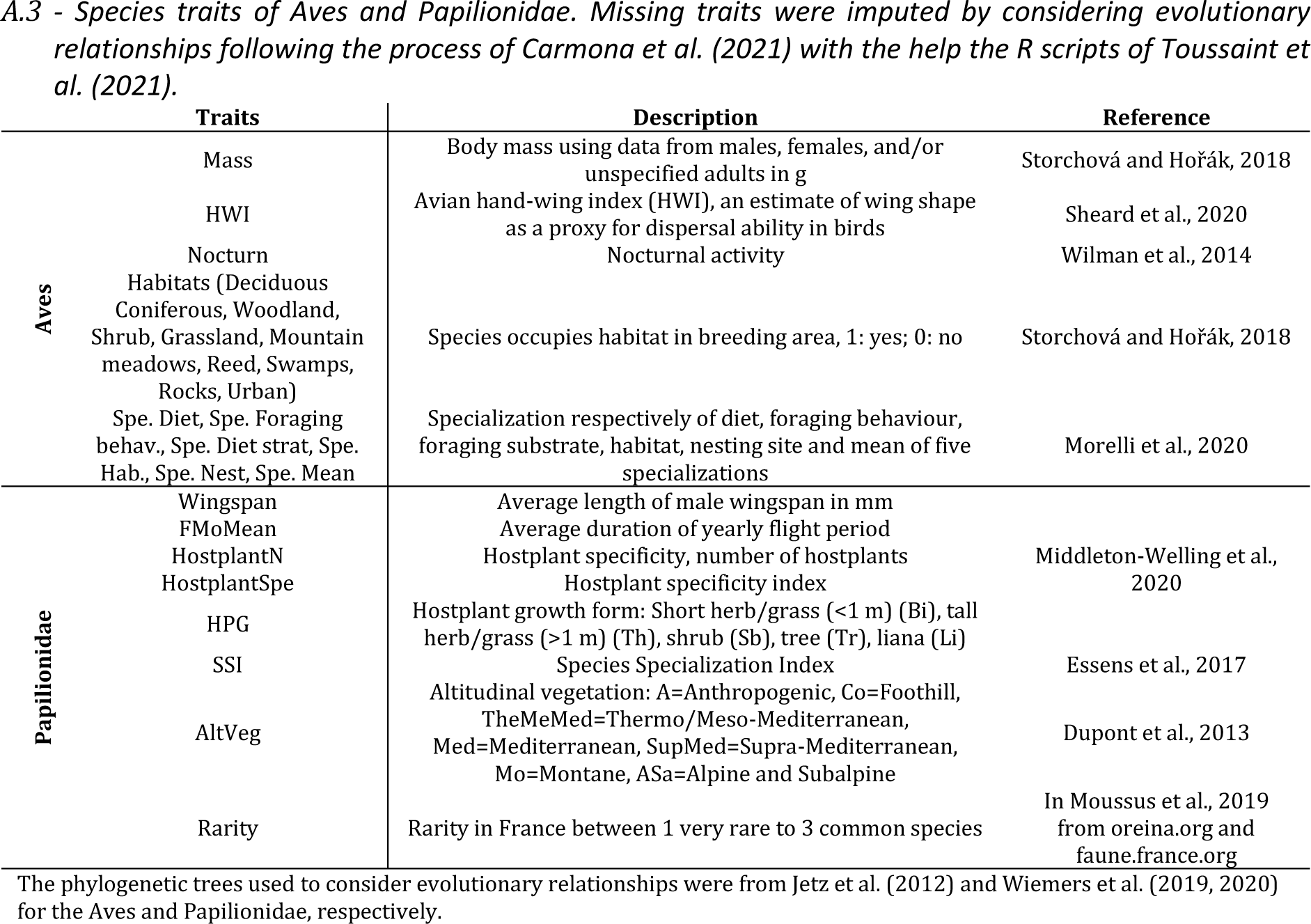

### Appendix B - Supplementary results

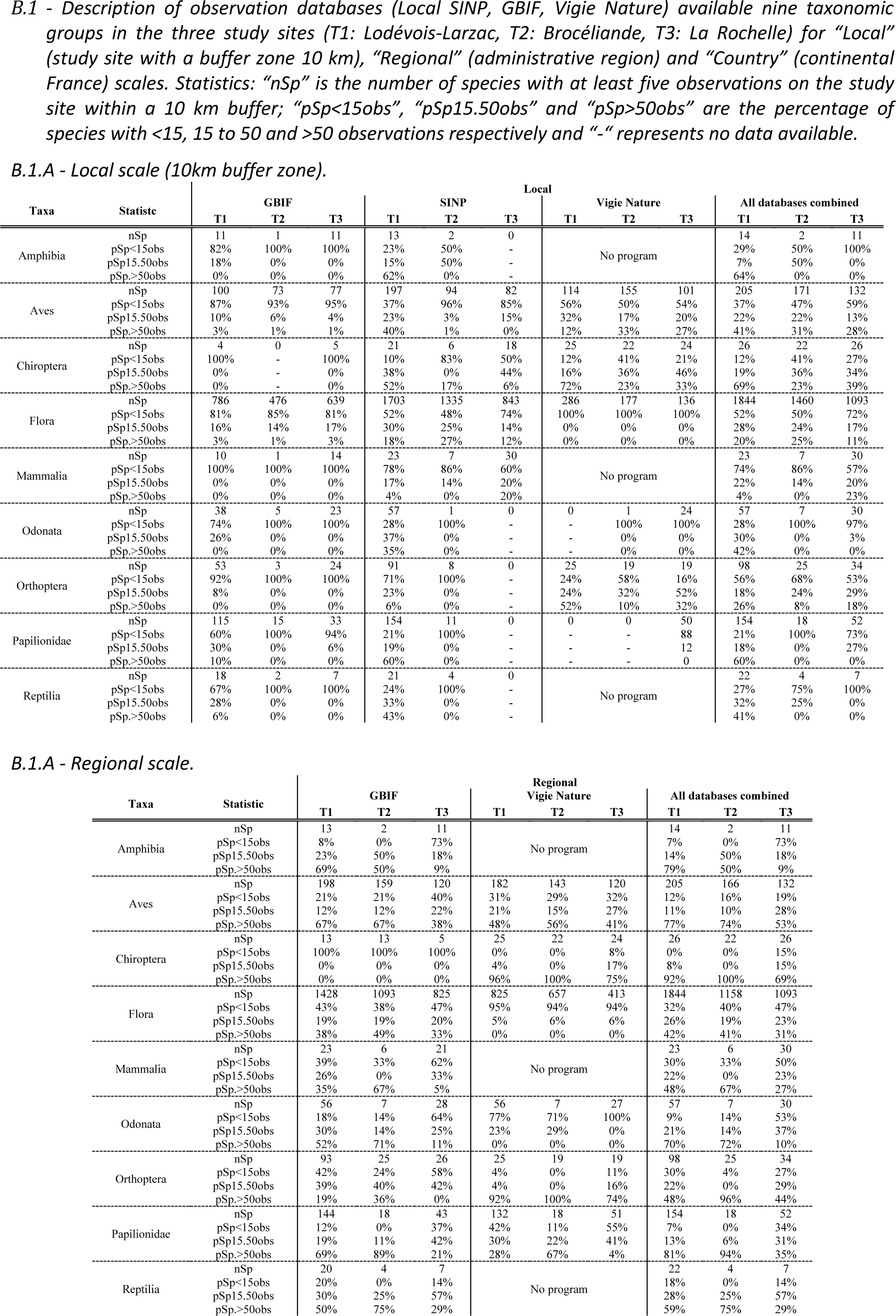
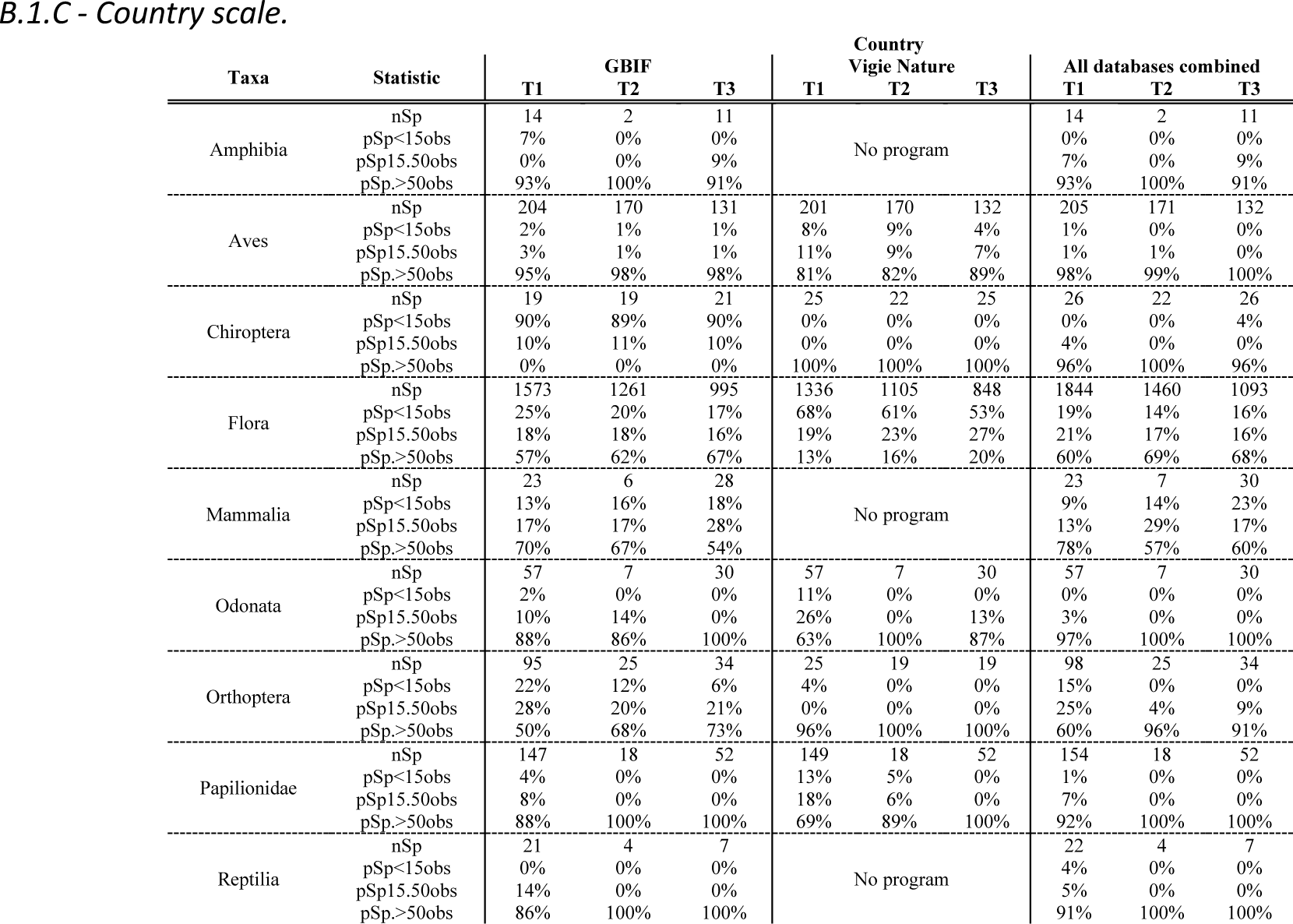
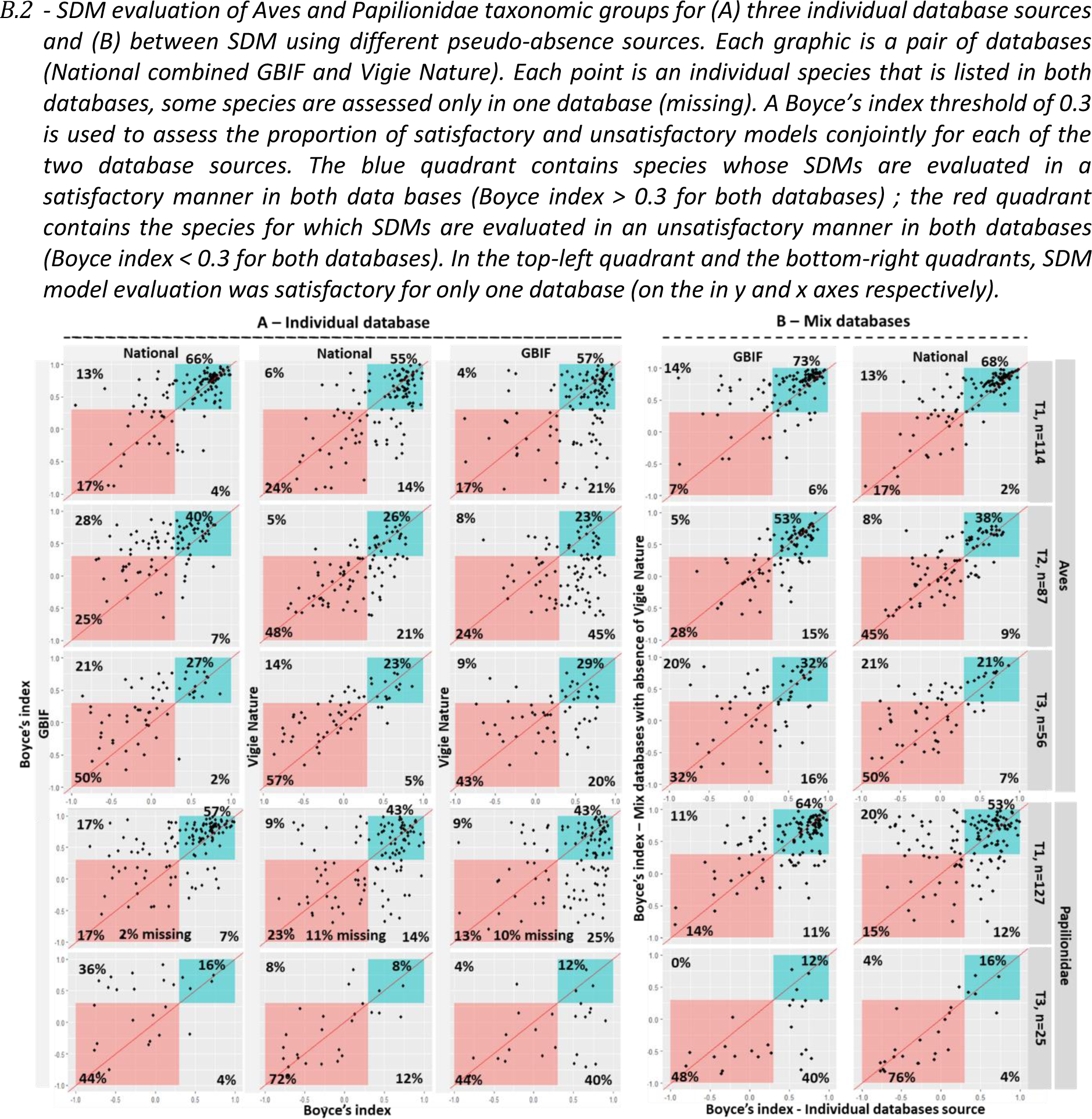
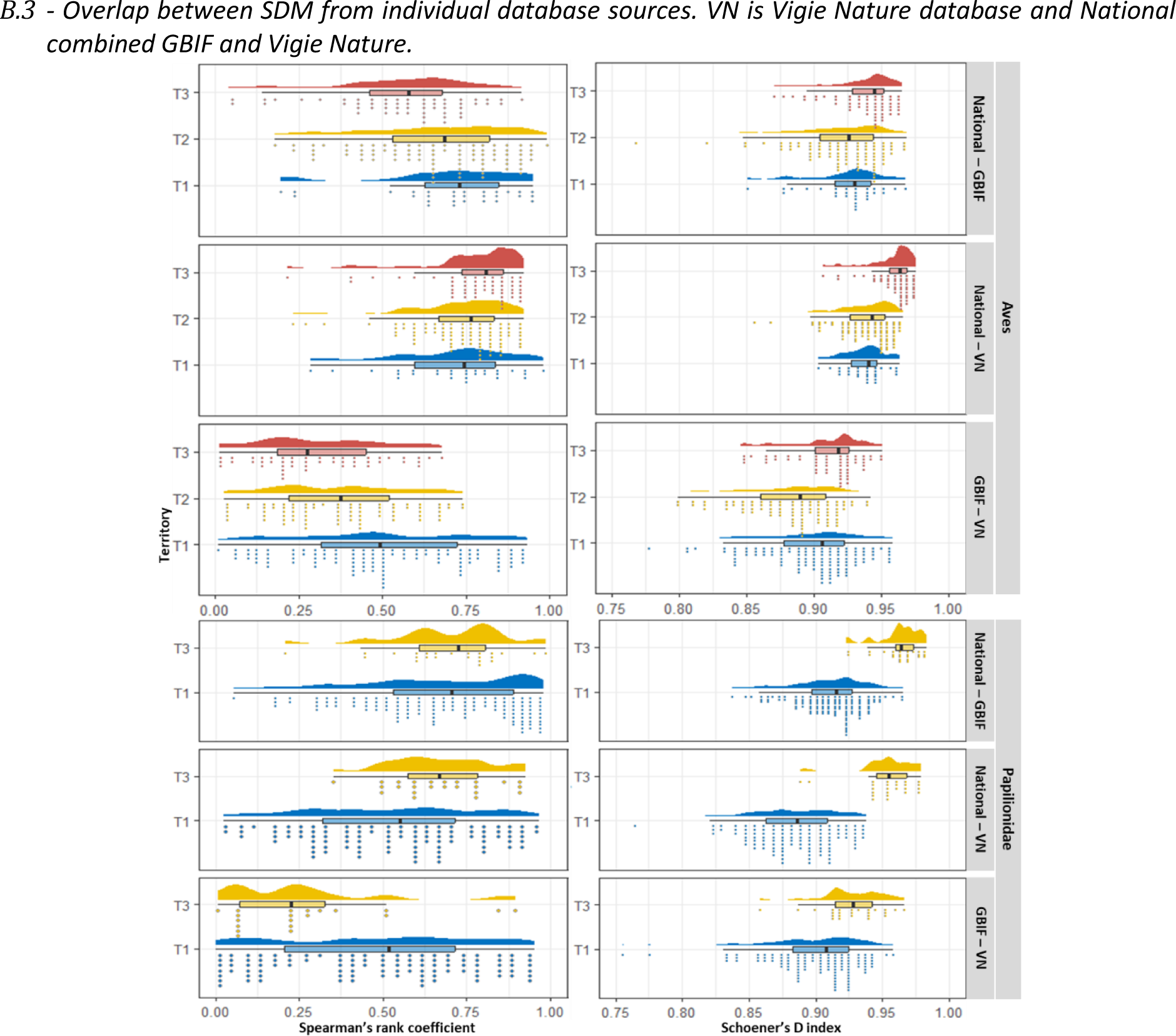
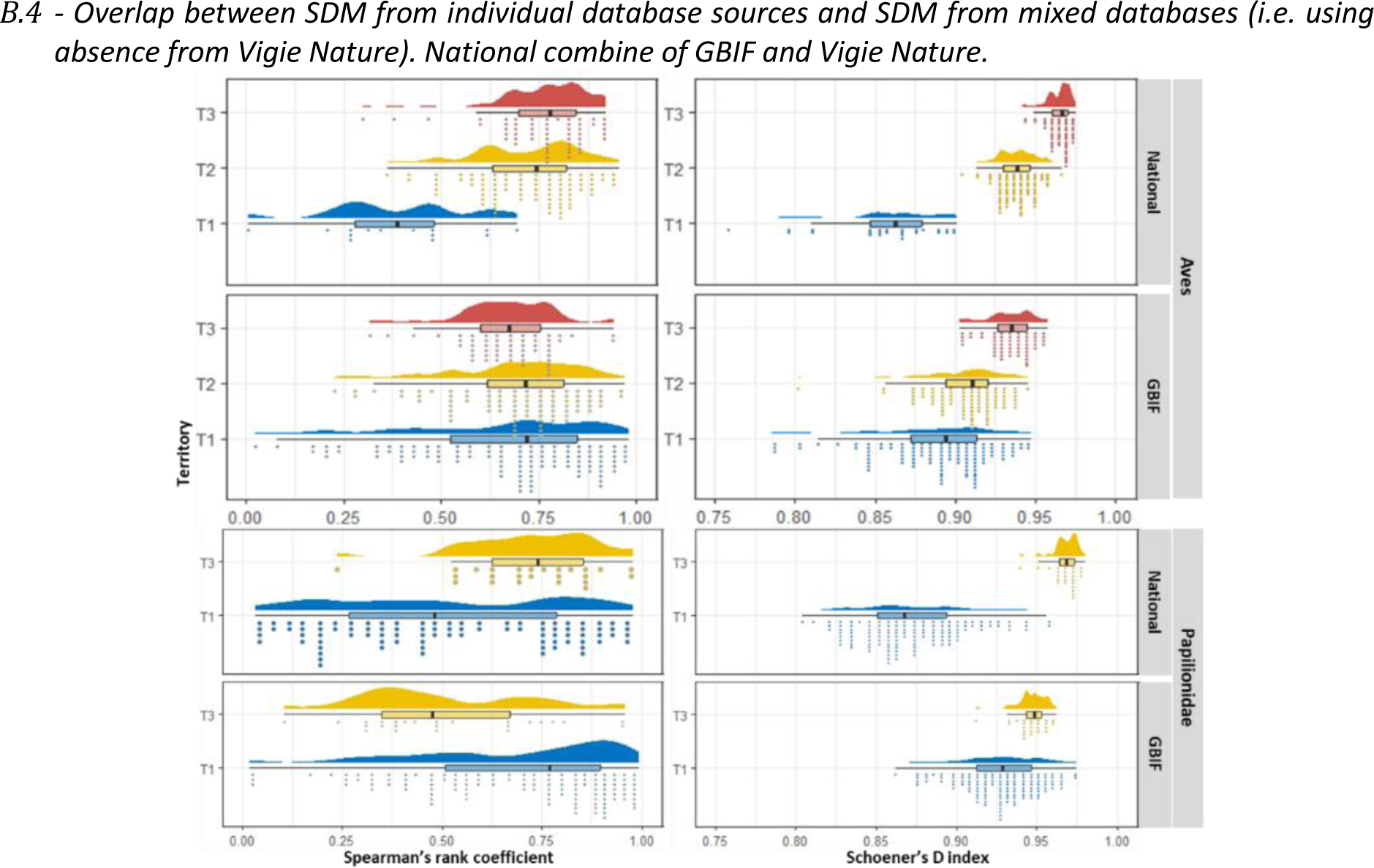
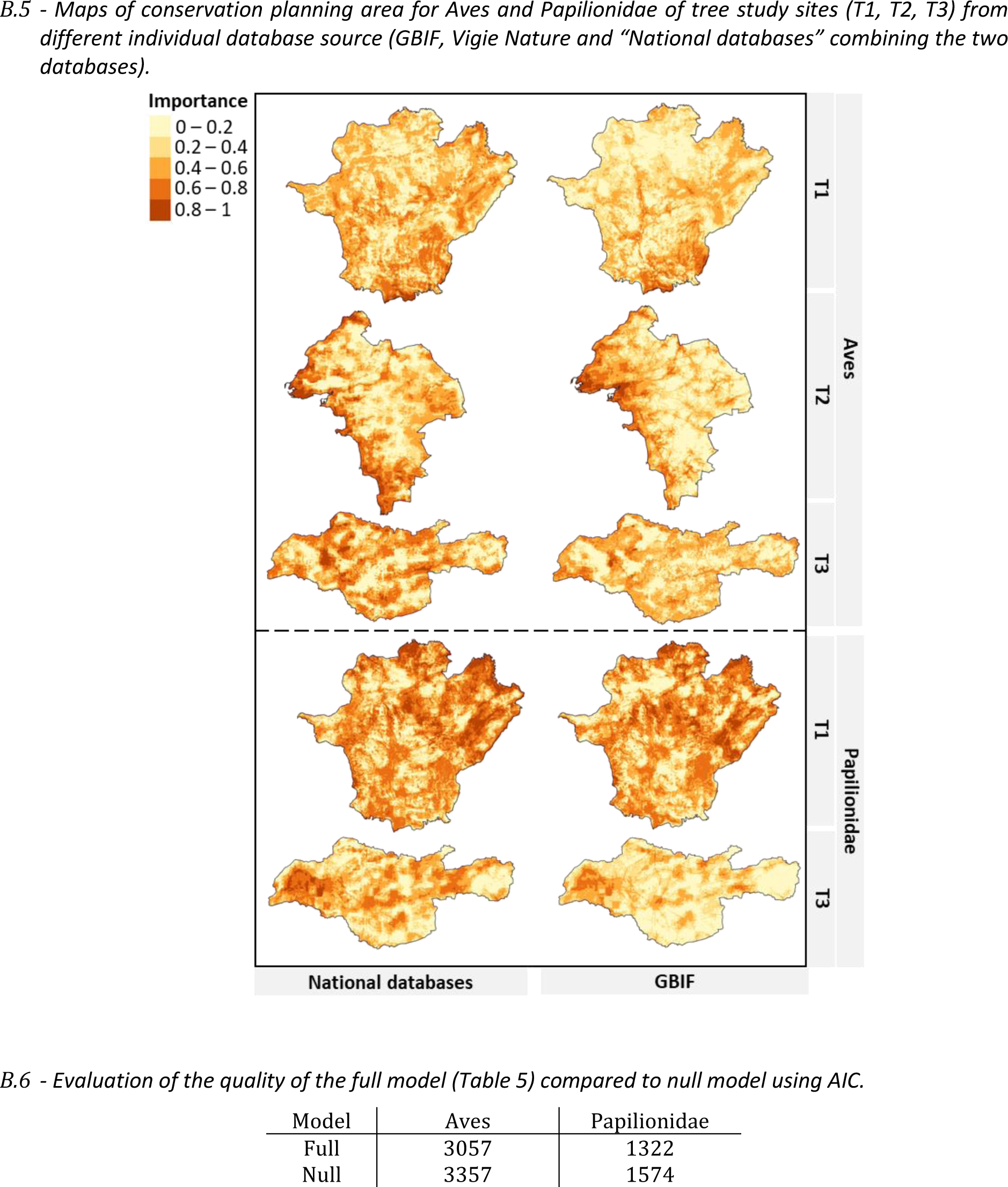
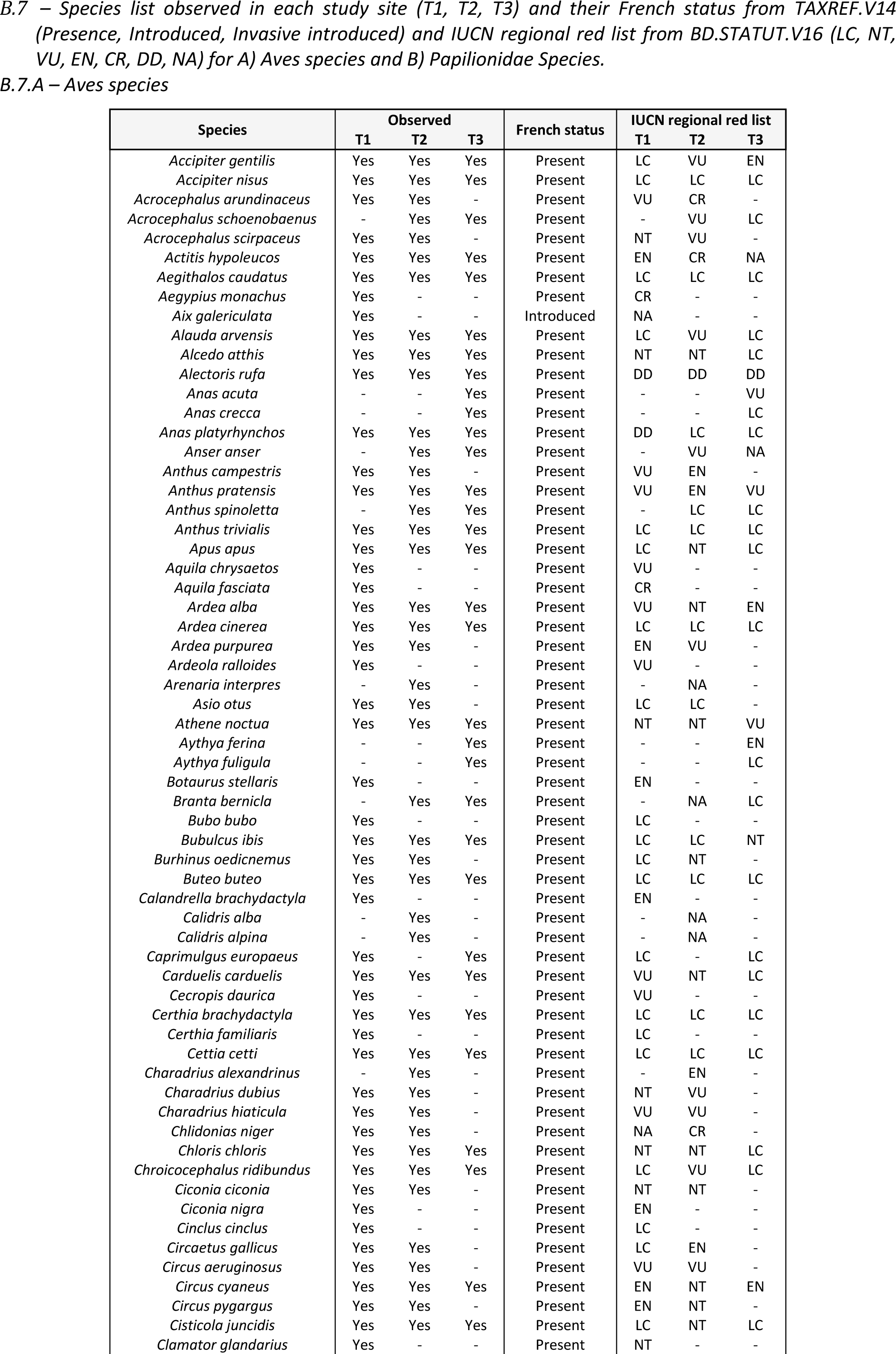
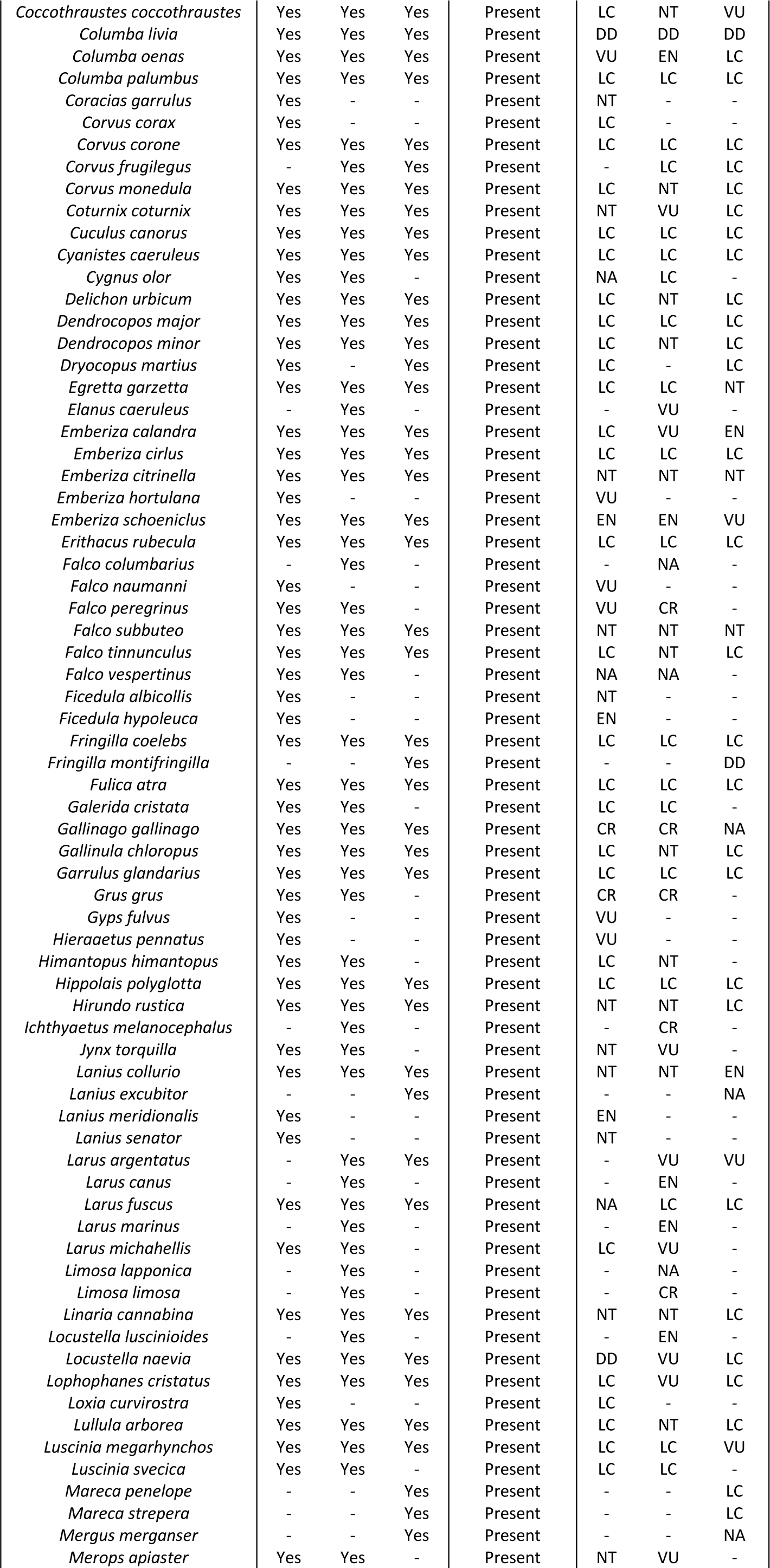
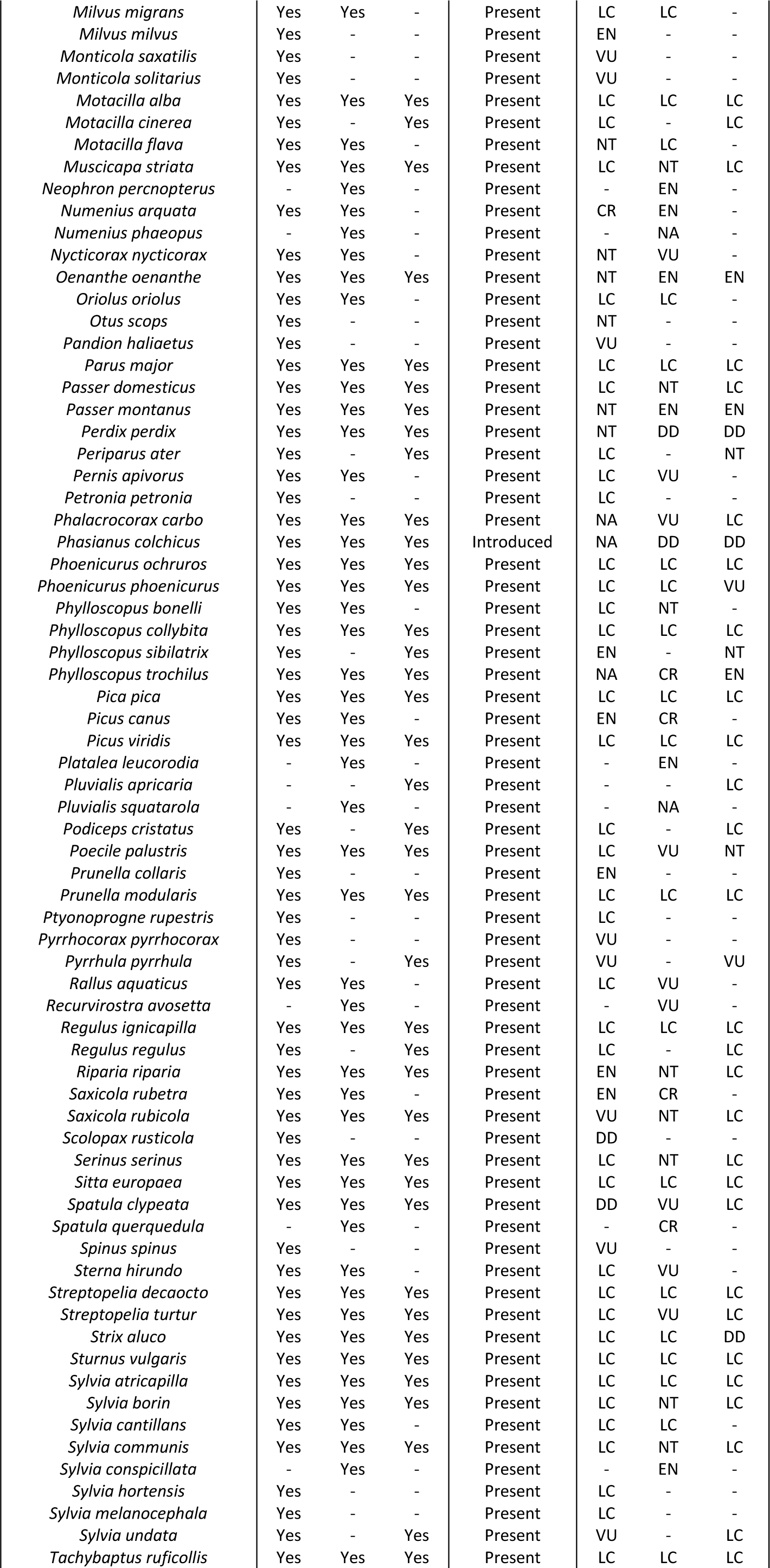
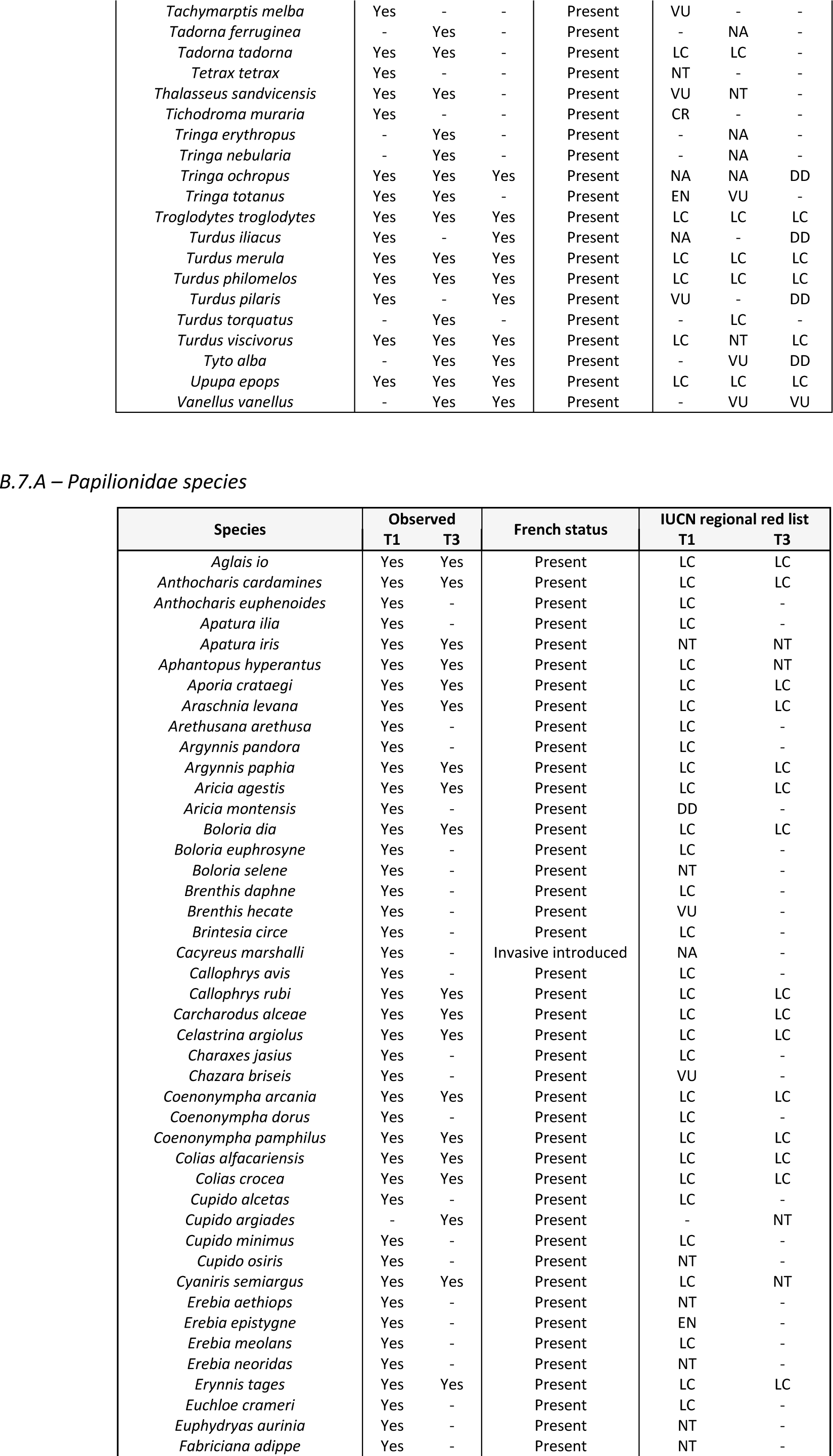
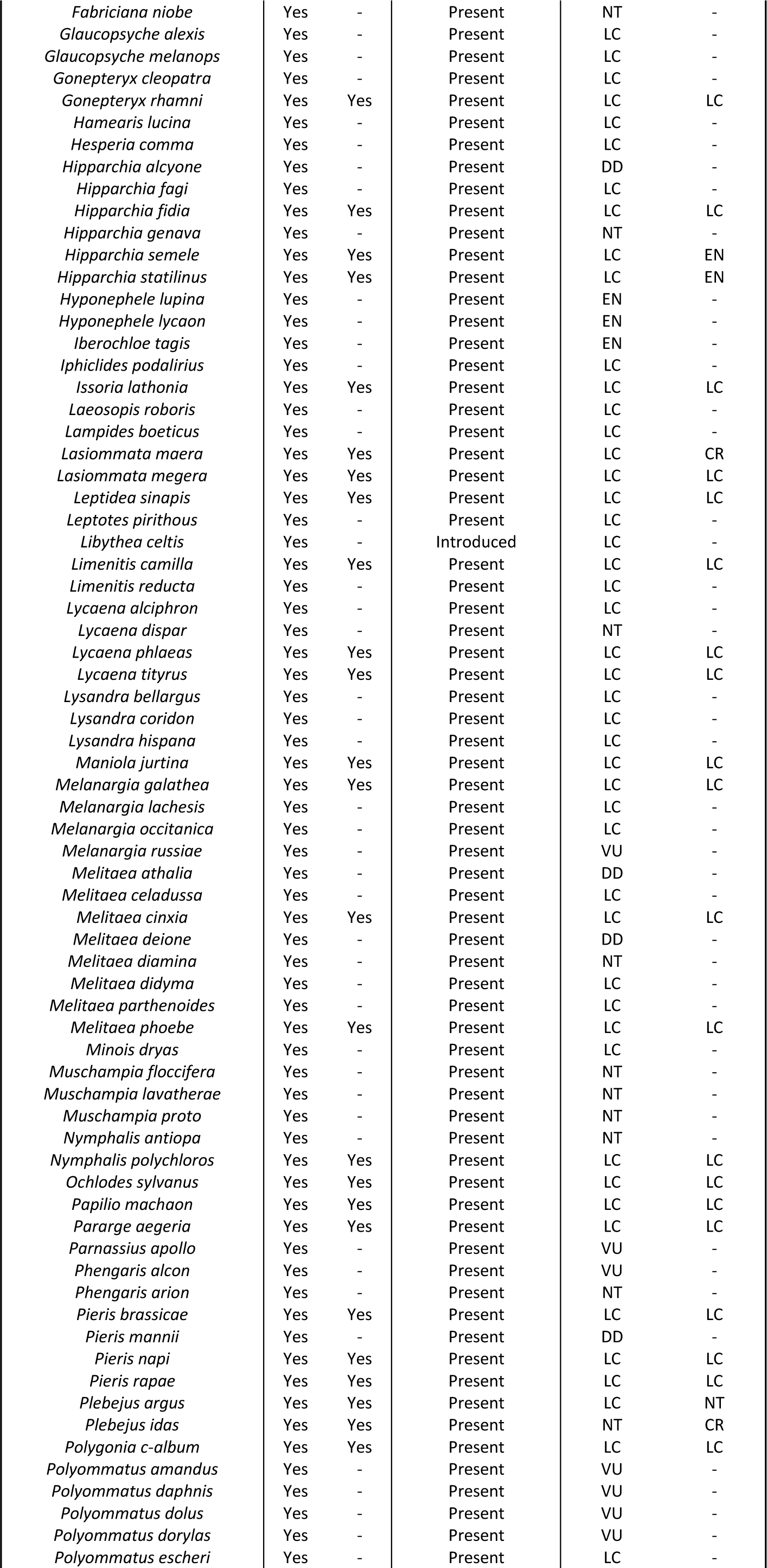
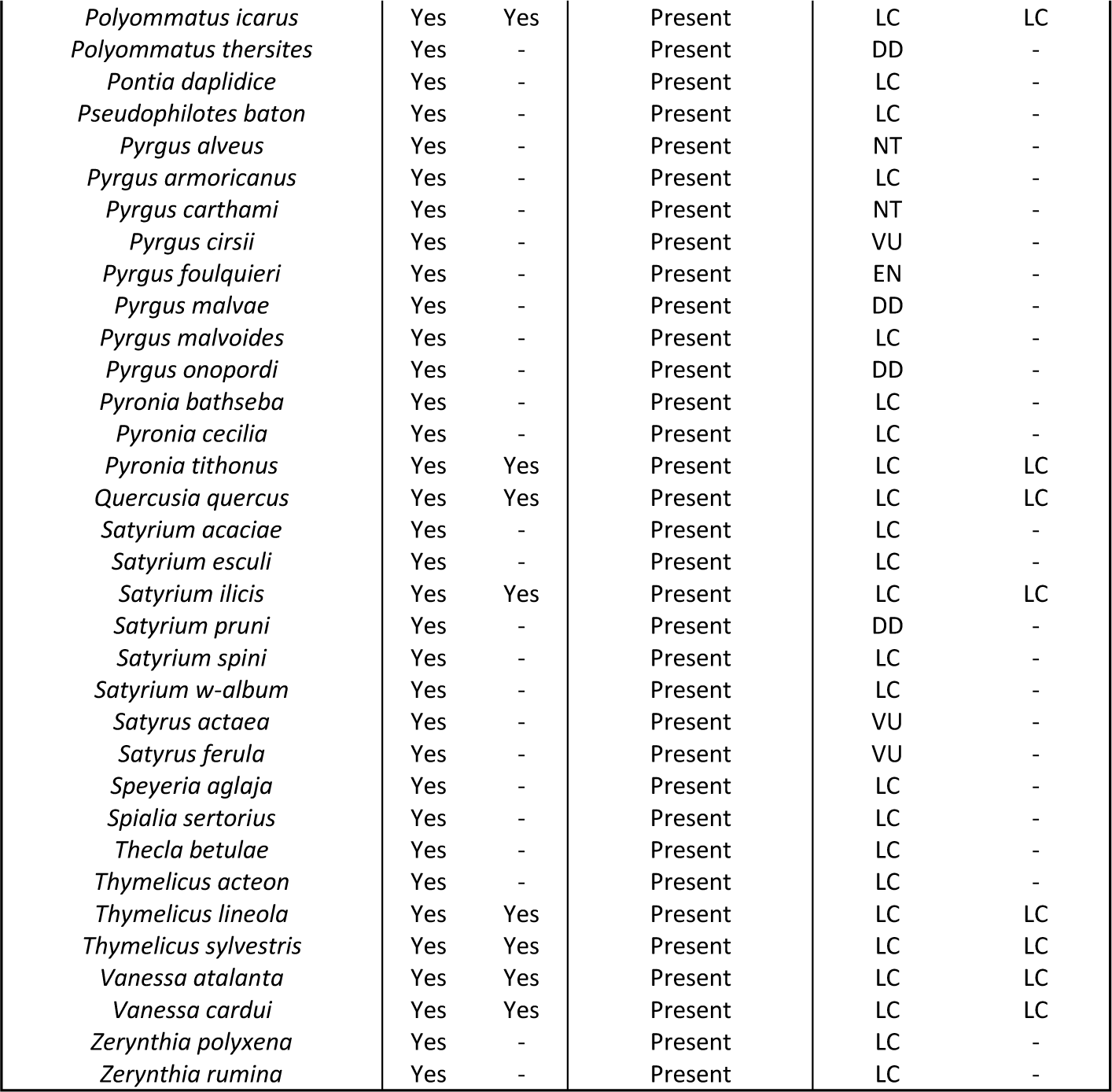

